# Broad sarbecovirus neutralizing antibodies define a key site of vulnerability on the SARS-CoV-2 spike protein

**DOI:** 10.1101/2020.05.15.096511

**Authors:** Anna Z. Wec, Daniel Wrapp, Andrew S. Herbert, Daniel Maurer, Denise Haslwanter, Mrunal Sakharkar, Rohit K. Jangra, M. Eugenia Dieterle, Asparouh Lilov, Deli Huang, Longping V. Tse, Nicole V. Johnson, Ching-Lin Hsieh, Nianshuang Wang, Juergen H. Nett, Elizabeth Champney, Irina Burnina, Michael Brown, Shu Lin, Melanie Sinclair, Carl Johnson, Sarat Pudi, Robert Bortz, Ariel S. Wirchnianski, Ethan Laudermilch, Catalina Florez, J. Maximilian Fels, Cecilia M. O’Brien, Barney S. Graham, David Nemazee, Dennis R. Burton, Ralph S. Baric, James E. Voss, Kartik Chandran, John M. Dye, Jason S. McLellan, Laura M. Walker

## Abstract

Broadly protective vaccines against known and pre-emergent coronaviruses are urgently needed. Critical to their development is a deeper understanding of cross-neutralizing antibody responses induced by natural human coronavirus (HCoV) infections. Here, we mined the memory B cell repertoire of a convalescent SARS donor and identified 200 SARS-CoV-2 binding antibodies that target multiple conserved sites on the spike (S) protein. A large proportion of the antibodies display high levels of somatic hypermutation and cross-react with circulating HCoVs, suggesting recall of pre-existing memory B cells (MBCs) elicited by prior HCoV infections. Several antibodies potently cross-neutralize SARS-CoV, SARS-CoV-2, and the bat SARS-like virus WIV1 by blocking receptor attachment and inducing S1 shedding. These antibodies represent promising candidates for therapeutic intervention and reveal a new target for the rational design of pan-sarbecovirus vaccines.

---

In December 2019, a novel pathogen emerged in the city of Wuhan in China’s Hubei province, causing an outbreak of atypical pneumonia (a disease known as COVID-19). The infectious agent was rapidly characterized as a lineage B betacoronavirus, named Severe Acute Respiratory Syndrome Coronavirus 2 (SARS-CoV-2) and shown to be closely related to SARS-CoV and several SARS-like bat CoVs(*1*). Despite the urgent need, there are currently no approved vaccines or therapeutics available for the prevention or treatment of COVID-19. Furthermore, the recurrent zoonotic spillover of CoVs into humans, along with the broad diversity of SARS-like CoV strains circulating in animal reservoirs, suggests that novel pathogenic CoVs are likely to emerge in the future and underscores the need for broadly active countermeasures.

CoV entry into host cells is mediated by the viral S glycoprotein, which forms trimeric spikes on the viral surface(*2*). Each monomer in the trimeric S assembly is a heterodimer of S1 and S2 subunits. The S1 subunit is composed of four domains: an N-terminal domain (NTD), a C-terminal domain (CTD), and subdomains I and II(*3*-*5*). The CTD of both SARS-CoV and SARS-CoV-2 functions as the receptor-binding domain (RBD) for the shared entry receptor, human angiotensin converting enzyme 2 (hACE2)(*6*-*10*). The S2 subunit contains the fusion peptide, heptad repeat 1 and 2, and a transmembrane domain, all of which are required for fusion of the viral and host cell membranes.

The S glycoprotein of human CoVs (HCoVs) is the primary target for neutralizing antibodies (nAbs)(*11*). Given that SARS-CoV and SARS-CoV-2 share about 80% amino acid identity in their S proteins, one important immunological question concerns the immunogenicity of conserved surfaces on these antigens. Studies of convalescent sera and a limited number of monoclonal antibodies (mAbs) have revealed limited to no cross-neutralizing activity, demonstrating that conserved antigenic sites are rarely targeted by nAbs(*5, 9, 12*). However, the frequencies, specificities, and functional activities of cross-reactive antibodies induced by natural SARS-CoV and SARS-CoV-2 infection remain poorly defined.

In this study, we aimed to comprehensively profile the cross-reactive B cell response induced by SARS-CoV infection by cloning an extensive panel of SARS-CoV-2 S-reactive mAbs from the peripheral B cells of a convalescent donor (Donor 84) who survived the 2003 SARS outbreak. To isolate cross-reactive antibodies, we stained purified B cells with a panel of memory B cell (MBC) markers and a fluorescently labeled recombinant SARS-CoV-2 S protein. Flow cytometric analysis revealed that 0.14% of class-switched MBCs were SARS-CoV-2 S-reactive, which was about 3-fold over background staining observed with a SARS-CoV-naïve donor sample (**Fig. 1A**). Notably, the frequency of antigen-specific MBCs was higher than expected, given the long interval between infection and blood draw (17 years) and previous studies showing waning of SARS-CoV-specific MBCs to undetectable levels within 6 years(*13*). Cognate antibody heavy- and light-chain pairs were rescued from 315 individual SARS-CoV-2-reactive B cells by single-cell RT-PCR and subsequently cloned and expressed as full-length IgGs in an engineered strain of *Saccharomyces cerevisiae(14)*. Of the 315 cloned antibodies, 200 bound to SARS-CoV-2 S in preliminary binding screens (**Fig. 1B**). Sequence analysis revealed that about half of the clones were members of expanded clonal lineages, whereas the other half were unique (**Fig. 1C**). This result is in stark contrast to numerous studies of other primary viral infections reporting very limited clonal expansion within virus-specific MBC repertoires(*15*-*18*). Moreover, about 30% of isolated antibodies displayed convergent VH1-69/VK2-30 germline gene pairing (**Fig. 1C**). As expected, almost all the antibodies were somatically mutated, with members of clonally expanded lineages showing significantly higher levels of somatic hypermutation (SHM) compared to unique clones (**Fig. 1D**). Finally, consistent with the respiratory nature of SARS-CoV infection, index sorting analysis revealed that 33% of binding antibodies originated from IgA^+^ MBCs and the remaining 66% from IgG^+^ MBCs (**Fig. 1E**). We conclude that SARS-CoV infection elicited a high frequency of long-lived, cross-reactive MBCs in this donor.

**Figure 1.**
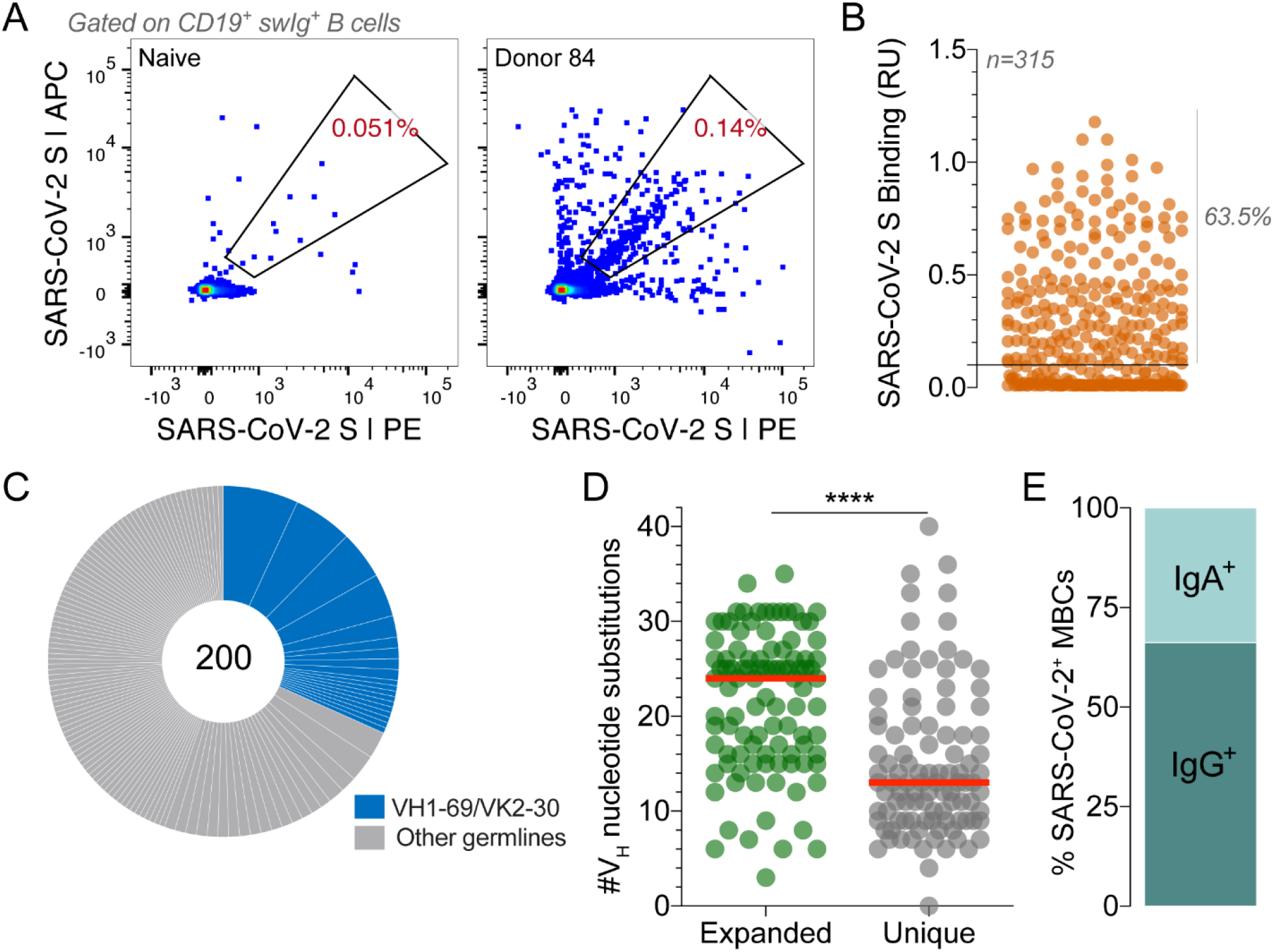
Isolation of SARS-CoV-2 S-specific IgGs. (**A**) Frequency of SARS-CoV-2 S-reactive B cells in Donor 84 and a negative control SARS-CoV-naïve donor. Fluorescence activated cell sorting (FACS) plots shown are gated on CD19^+^CD20^+^IgD^—^IgM^—^ B cells. SARS-CoV-2 S was labeled with two different colors to reduce background binding. The percentage shown in the gate indicates the frequency of SARS-CoV-2 S-reactive B cells among CD19^+^CD20^+^IgD^—^IgM^—^ B cells. (**B**) Binding of 315 isolated antibodies to SARS-CoV-2 S, as determined by biolayer interferometry (BLI). The solid line indicates the threshold used for designating binders (0.1 RUs). (**C**) Clonal lineage analysis. Each lineage is represented as a segment proportional to the lineage size. Clones that that utilize VH1-69/VK2-30 germline gene pairing are shown in blue. The total number of isolated antibodies is shown in the center of the pie. Clonal lineages were defined based on the following criteria: identical VH and VL germline genes, identical CDR H3 length, and CDR H3 amino acid identity ≥80%. (**D**) Load of somatic mutations, expressed as number of nucleotide substitutions in VH, in unique antibodies and members of expanded clonal lineages. (**E**) Proportion of SARS-CoV-2 S binding antibodies derived from IgG^+^ and IgA^+^ B cells, as determined by index sorting. Statistical comparisons were made using the Mann-Whitney test (**** P < 0.0001). Red bars indicate medians. swIg, switched immunoglobulin; RU, response units; VH, variable region of the heavy chain.

We next measured the apparent binding affinities (K_D_^Apps^) of the antibodies to prefusion-stabilized SARS-CoV and SARS-CoV-2 S proteins(*5*). Although the majority of antibodies (153 out of 200) showed binding to both S proteins, a subset appeared to be SARS-CoV-2 S-specific (**Fig. 2A**). This result was unexpected given that the antibodies were isolated from a SARS-CoV-experienced donor and may relate to differences between the infecting SARS-CoV strain and the recombinant SARS-CoV S protein (Tor2) used for the binding studies. Alternatively, this result may be due to inherent differences in the stability or antigenicity of recombinant prefusion-stabilized SARS-CoV and SARS-CoV-2 S proteins. Indeed, about 30% of antibodies that failed to bind recombinant SARS-CoV S displayed reactivity with SARS-CoV S expressed on the surface of transfected cells, providing some evidence for differences in the antigenicity of recombinant and cell-expressed forms of S **(Fig. S1**).

**Figure 2.**
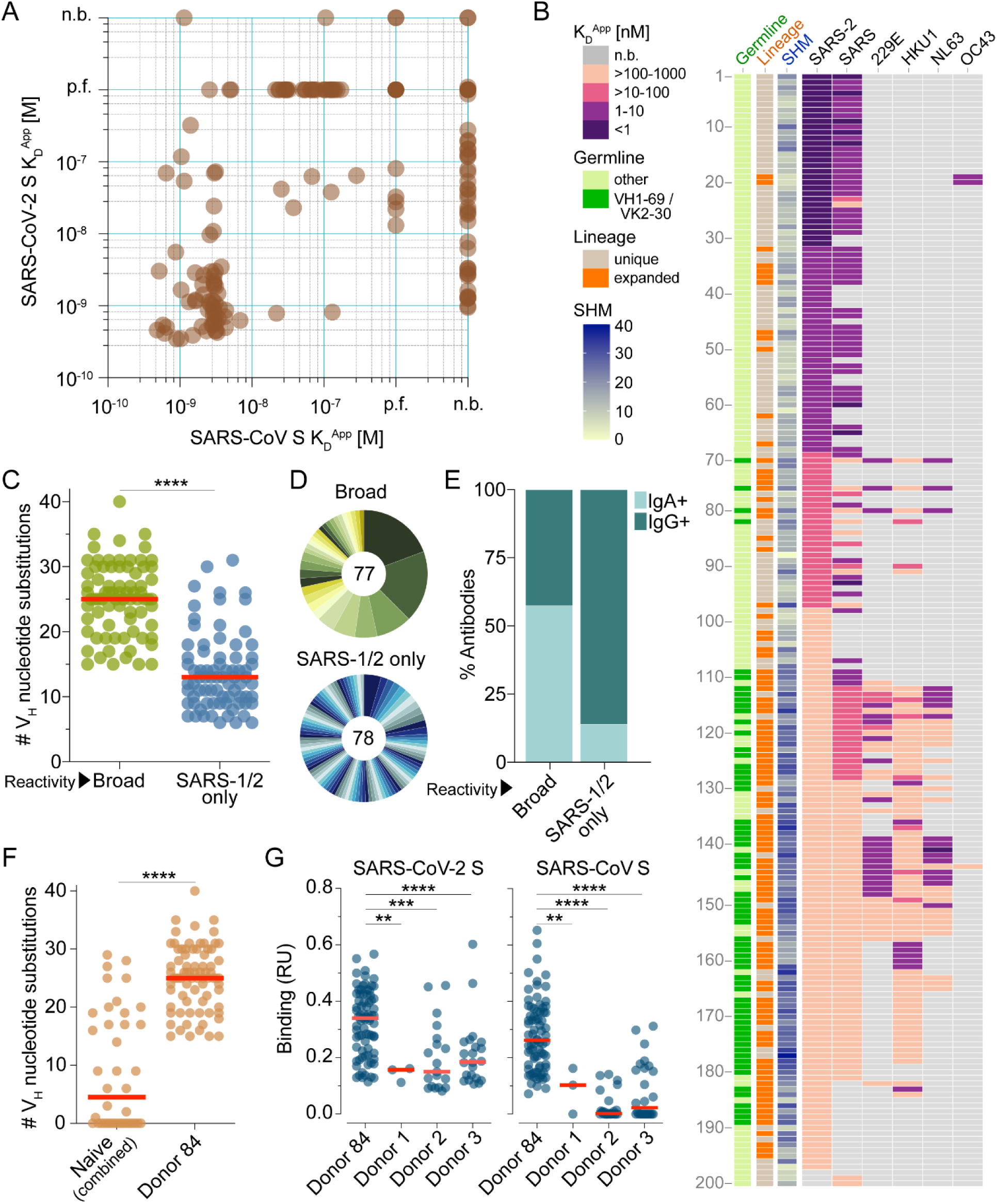
Binding properties of SARS-CoV-2 S-specific antibodies. (**A**) Apparent binding affinities of SARS-CoV-2 S-specific IgGs for prefusion-stabilized SARS-CoV and SARS-CoV-2 S proteins, as determined by BLI measurements. Low affinity clones for which binding curves could not be fit are designated as “poor fit” on the plot. (**B**) Apparent binding affinities of the isolated antibodies for SARS-CoV-2, SARS-CoV, 229E, HKU1, NL63, and OC43 S proteins. Germline gene usage, clonal expansion, and SHM are indicated in the three leftmost panels. SHM is represented as the number of nucleotide substitutions in VH. (**C**) Load of somatic mutations in broadly cross-reactive and SARS-CoV/SARS-CoV-2-specific antibodies. Red bars indicate medians. (**D**) Degree of clonal expansion in broadly cross-reactive and SARS-CoV/SARS-CoV-2-specific antibodies. Each lineage is represented as a segment proportional to the lineage size. The total number of antibodies is shown in the center of the pie. (**E**) Proportion of broadly cross-reactive and SARS-CoV/SARS-CoV-2-specific antibodies derived from IgG^+^ and IgA^+^ B cells, as determined by index sorting. **(F**) Load of somatic mutations in SARS-CoV-2 S-reactive antibodies isolated from three naive donors and Donor 84. Antibodies from naïve donors were combined for this analysis. (**G**) Binding activity of antibodies isolated from Donor 84 and three naïve donors to SARS-CoV and SARS-CoV-2 S, as determined by BLI. p.f., poor fit; n.b., non-binder; RU, response units. Statistical comparisons were made using the Mann-Whitney test (** P < 0.01; *** P < 0.001; **** P < 0.0001).

Paradoxically, most of the highly mutated and clonally expanded antibodies bound weakly (K_D_^Apps^ >10 nM) to both SARS-CoV and SARS-CoV-2 S (**Fig. 2B**). We sought to determine if these antibodies originated from pre-existing MBCs that were induced by prior exposures to naturally circulating HCoVs, which share up to 32% amino acid identity with SARS-CoV and SARS-CoV-2 in their S proteins. Accordingly, we assessed binding of the antibodies to recombinant S proteins of naturally circulating human alphacoronaviruses (HCoV-NL63 and HCoV-229E) and betacoronaviruses (HCoV-OC43 and HCoV-HKU1). Over 80% of the low affinity (K_D_^Apps^ >10 nM) SARS-CoV/SARS-CoV-2 cross-reactive antibodies showed reactivity with one or more of the HCoV S proteins, suggesting SARS-CoV infection may have boosted a pre-existing MBC response induced by circulating HCoVs (**Fig. 2B**). Alternatively, circulating HCoV infections experienced by this donor after the SARS-CoV infection may have expanded this cross-reactive MBC pool. Consistent with this hypothesis, the broadly cross-reactive antibodies showed significantly higher levels of SHM and clonal expansion compared to those that only recognized SARS-CoV and SARS-CoV-2 (**Fig. 2B-D**). Furthermore, 72% of the broadly binding antibodies utilized VH1-69/VK2-30 germline gene pairing, suggesting germline-mediated recognition of a common antigenic site (**Fig. 2B** and **Fig. S2**). Index sorting analysis revealed that the majority of the broadly cross-reactive antibodies were derived from IgA^+^ MBCs, indicating a mucosal origin, whereas most of the SARS-CoV/SARS-CoV-2 cross-reactive antibodies originated from IgG^+^ MBCs (**Fig. 2E**). Finally, the vast majority of cross-reactive antibodies lacked polyreactivity, demonstrating that their broad binding activity is not due to non-specific cross-reactivity (**Fig. S3**).

To investigate whether the above results were due to an “original antigenic sin” (OAS) phenomenon, or rather simply due to avid binding of circulating HCoV-specific B cell receptors to the SARS-CoV-2 S tetramers used for cell sorting, we assessed whether similarly broadly binding antibodies were also present in SARS-CoV/SARS-CoV-2-naïve donors that had been exposed to endemic HCoVs. We obtained peripheral blood mononuclear cell (PBMC) samples from three healthy adult donors with serological evidence of circulating HCoV exposure and no history of SARS-CoV or SARS-CoV-2 infection, and stained the corresponding B cells with a fluorescently labeled SARS-CoV-2 S probe (**Fig. S4A**). Flow cytometric analysis revealed that between 0.06-0.12% of total B cells in the three naïve donors displayed SARS-CoV-2 reactivity (**Fig. S4B**). Over 350 SARS-CoV-2-reactive MBCs were sorted and amplified by single-cell RT-PCR, and 141 V_H_/V_L_ pairs were cloned and expressed as full-length IgGs. Although a limited number of SARS-CoV-2 S binding antibodies were isolated from all three naïve donors, they displayed significantly lower levels of SHM, clonal expansion, and binding affinities for both SARS-CoV and SARS-CoV-2 S compared to the cross-reactive antibodies identified from the convalescent SARS donor **(Fig. 2F-G** and **Fig. S4C**). Altogether, these results suggest that SARS-CoV infection likely led to the activation and expansion of pre-existing cross-reactive MBCs induced by circulating HCoV exposure in this donor.

To map the antigenic sites recognized by the SARS-CoV/SARS-CoV-2 cross-reactive antibodies, we performed a series of binding experiments using a panel of recombinant S protein subunits and individual domains. Due to the inherent technical challenges associated with measuring binding of low affinity antibodies to monomeric proteins, we analyzed only the 64 high affinity binders (K_D_^Apps^<10 nM) to SARS-CoV-2 S (**Fig. 2A, B**). We first evaluated binding to recombinant SARS-CoV-2 S1 and S2 subunits and observed that 75% of the antibodies recognized epitopes within S1, whereas the remaining 25% bound to epitopes within S2 (**Fig. 3A**). Interestingly, two of the S2-directed antibodies also showed strong reactivity with OC43 S, suggesting recognition of a highly conserved antigenic site (**Fig. S5**). We next evaluated the 49 S1-directed antibodies for reactivity with individual SARS-CoV-2 RBD and NTD domains and found that 21 (43%) and 28 (57%) of the S1-specific antibodies recognized the RBD and NTD, respectively (**Fig. 3A**).

**Figure 3.**
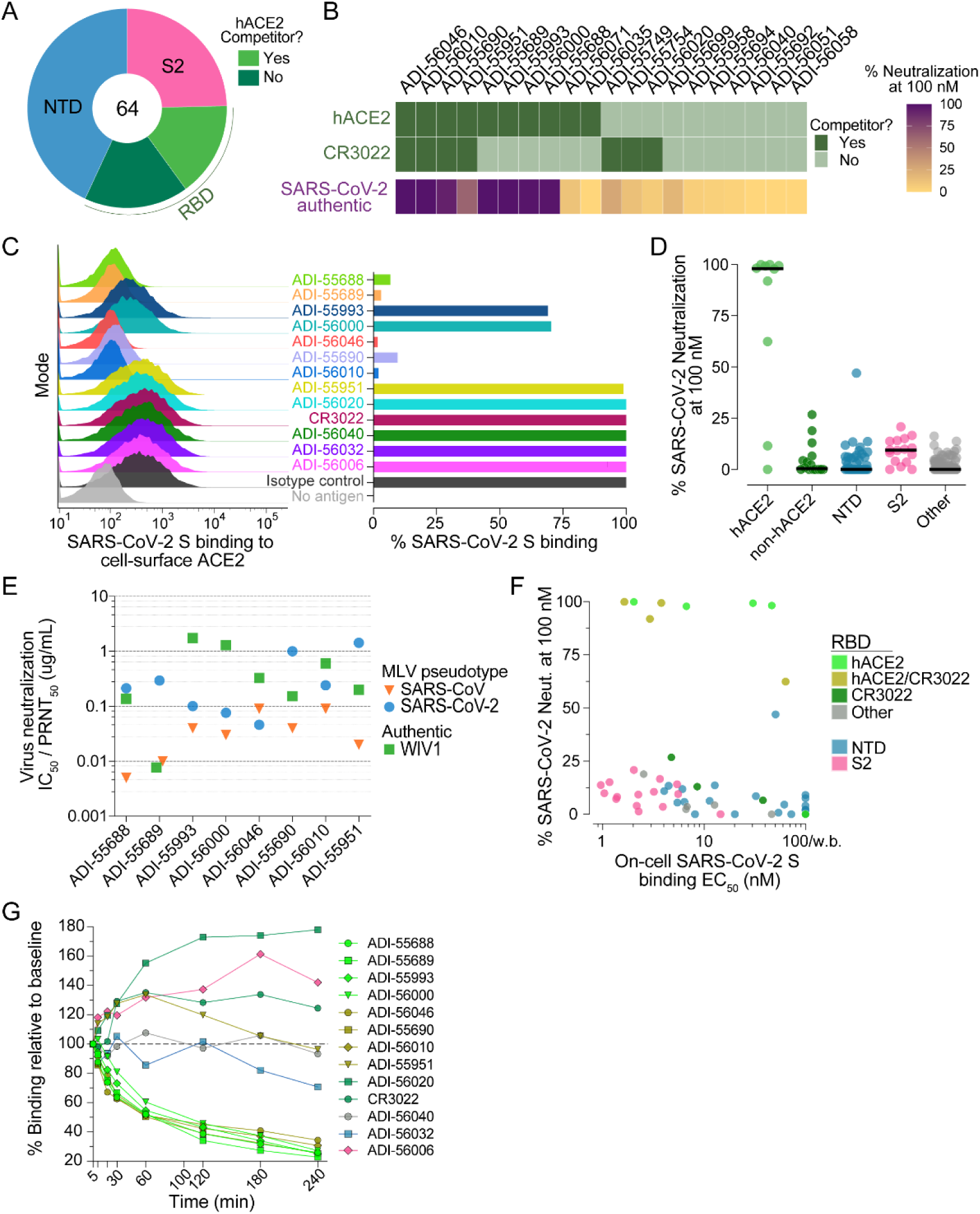
Epitope mapping and neutralization screening. **(A)** Proportion of SARS-CoV-2 S-specific antibodies targeting each of the indicated antigenic sites. **(B)** Heat map showing the competitive binding profiles of the RBD-directed antibodies (top) and percent neutralization of authentic SARS-CoV-2 at a 100 nM concentration (bottom). **(C)** Antibody inhibition of SARS-CoV-2 S binding to endogenous ACE2 expressed on Vero E6 cells, as determined by flow cytometry. Antibodies were mixed with recombinant SARS-CoV-2 S expressing a Twin-Strep-tag at a molar ratio of 5:1 before adding to Vero E6 cells. Strep-Tactin-PE was used to detect the relative intensity of SARS-CoV-2 S binding to cell-surface ACE2. An anti-ebolavirus antibody (KZ52) was used as an isotype control. The “no antigen” control shown in the right panel indicates secondary-only staining. Percent binding shown in the right panel was normalized to isotype control. **(D)** Percent authentic SARS-CoV-2 neutralization observed in the presence of 100 nM antibody. Antibodies are grouped according to epitope specificity. RBD-directed antibodies that compete or do not compete with ACE2 are designed as ACE2 and non-ACE2, respectively. **(E)** Antibody neutralization of SARS-CoV and SARS-CoV-2 MLV pseudovirus (strain n-CoV/USA_WA1/2020) using HeLa-ACE2 target cells, and antibody neutralization of authentic WIV1-CoV using Vero E6 target cells. SARS-CoV and SARS-CoV-2 IC_50_s and WIV1-CoV PRNT_50_s are reported in µg/ml. **(F)** Binding EC_50_s for cell-surface SARS-CoV-2 S are plotted against percent neutralization of authentic SARS-CoV-2 at 100 nM. Background binding was assessed using mock transfected HEK293 cells. Data points are colored according to epitope specificity. RBD-directed antibodies are further categorized based on their competition group: hACE2, antibodies that only compete with hACE2; CR3022, antibodies that only compete with CR3022; hACE2/CR3022, antibodies that compete with hACE2 and CR3022; Other, antibodies that do not compete with hACE2 or CR3022. **(G)** Antibody binding activity to cell-surface SARS-CoV-2 S over time, as determined by flow cytometry. IgGs were incubated with cells expressing WT SARS-CoV-2 at 37°C and aliquots were placed on ice at the indicated time points. Binding MFI was assessed at 240 min for all samples. CR3022 is included for comparison. Curves are colored by epitope specificity, as in (F).

To further define the epitopes recognized by the RBD-directed antibodies, we performed competitive binding studies with recombinant hACE2 and a previously described antibody, CR3022, that targets a conserved epitope that is distinct from the receptor binding site (**Fig. 3B** and **Fig. S6A**)(*19*). Six of the antibodies competed only with hACE2, three competed only with CR3022, four competed with both hACE2 and CR3022, and seven did not compete with hACE2 or CR3022 (**Fig. 3B**). Thus, these antibodies delineate at least four adjacent and potentially overlapping sites within the RBD. Importantly, antibodies that competed with recombinant hACE2 binding to SARS-CoV-2 RBD in the biolayer interferometry assay also interfered with binding of full-length SARS-CoV-2 S to endogenous ACE2 expressed on the surface of Vero E6 cells (**Fig. 3C**). In summary, SARS-CoV infection elicited high affinity cross-reactive antibodies to a range of antigenic sites within both the S1 and S2 subunits.

To evaluate the neutralization activities of the SARS-CoV-2 binding antibodies, we performed neutralization assays using both murine leukemia virus (MLV)- and vesicular stomatitis virus (VSV)-based pseudotype systems as well as authentic SARS-CoV-2. Due to the large number of antibodies, initial neutralization screening was performed in the authentic virus assay using a single concentration of purified IgG. Only nine out of 200 antibodies displayed neutralizing activity at the 100 nM concentration tested, eight of which targeted the RBD and the remaining one recognized the NTD (**Fig. 3D**). Similar results were observed in the VSV-based pseudovirus assay (**Fig. S7**). Of the eight RBD-directed nAbs, four targeted epitopes overlapping with both hACE2 and CR3022 and the other four recognized epitopes only overlapping that of hACE2, suggesting the existence of two partially overlapping neutralizing epitopes within the RBD (**Fig. 3B**). Neutralization titration studies revealed that the half maximal inhibitory concentrations (IC_50_s) of the RBD-directed nAbs ranged from 0.05-1.4 µg/ml against SARS-CoV-2 and 0.004-0.06 µg/ml against SARS-CoV in the MLV assay (**Fig. 3E** and **Fig. S8**). The SARS-CoV-2 neutralization IC_50_s observed in the MLV assay were within 1.5- to 5-fold of those observed in the authentic virus neutralization assay (**Fig. S8 and Fig. S9**). In contrast, the VSV-SARS-CoV-2 neutralization potencies were substantially elevated (8- to 35-fold) compared to those observed for live SARS-CoV-2, highlighting the need for standardized pseudovirus neutralization assays for the assessment of nAbs against SARS-CoV-2 (**Fig. S8 and Fig. S9**). To determine breadth of neutralization against representative pre-emergent SARS-like bat CoVs, we also carried out neutralization assays against WIV1-CoV using a classic plaque reduction assay(*20*). All eight antibodies neutralized WIV1-CoV with PRNT_50_s ranging from 0.076-1.7 µg/ml, demonstrating their breadth of activity (**Fig. 3E** and **Fig. S10**). Crucially, none of the antibodies left an un-neutralized viral fraction in any of the assays (**Fig. S8 and Fig. S10**).

Unexpectedly, we observed little to no correlation between binding affinity for WT SARS-CoV-2 cell surface S and neutralizing activity. For example, all of the S2-directed antibodies and a subset of NTD-directed antibodies bound with high affinity to both recombinant and cell surface S, but none were neutralizing (**Fig. 3F**). Similarly, the RBD-directed antibodies targeting epitopes outside of the hACE2 binding site showed little to no neutralizing activity, despite binding with similar affinity to cell surface S compared to many of the hACE2 competitor nAbs (**Fig. 3F** and **Fig. S11**). Although the reason(s) for the discrepancy between binding and neutralization are not yet clear, it is possible that the high affinity non-neutralizing antibodies recognize non-functional forms of S presented on the viral surface, as observed for many HIV non-neutralizing antibodies(*21*). Additionally, dissociation of the S1 subunit following receptor engagement may impede the ability of non-receptor blocking RBD- and NTD-directed antibodies to neutralize infection(*22*). Surprisingly, even within the group of hACE2-blocking nAbs, we did not observe a strong correlation between binding to cell surface-or recombinant-S and neutralizing activity, suggesting that antibody potency is governed at least in part by factors beyond binding affinity (**Fig. 3F** and **Fig. S12**). To determine whether the hACE2 competitor antibodies neutralized by inducing S1 shedding and premature S triggering(*23*), we incubated HEK-293 cells expressing WT SARS-CoV-2 S with saturating concentrations of antibody and measured the median fluorescence intensity (MFI) of antibody binding over time by flow cytometry. Indeed, all of the hACE2 blocking antibodies showed significantly decreased binding over time, consistent with induced S1 dissociation, whereas antibodies recognizing the NTD, S2 stem, and RBD epitopes outside of the hACE2 binding site displayed either no change or an increase in binding over the course of the experiment (240 minutes) (**Fig. 3G**). We conclude that SARS-CoV infection induces high affinity cross-reactive antibodies to multiple distinct antigenic sites on the S protein, but neutralizing activity is primarily restricted to RBD-directed antibodies that interfere with receptor binding and promote S1 dissociation.

To structurally characterize the epitopes recognized by the RBD-directed nAbs, we performed negative stain electron microscopy (EM) to observe each of these Fabs bound to the SARS-CoV-2 S protein. Many of the 2D class averages that we obtained displayed obvious heterogeneity in the number Fabs that were bound to a single S trimer, which is likely due to dynamic inaccessibility of RBD epitopes and sub-stoichiometric binding of S at the low protein concentrations used to prepare grids (**Fig. 4A**)(*5, 24*). The 3D reconstructions of these complexes support the results of our biophysical competition assays and show that the RBD-directed nAbs recognize an overlapping patch on the solvent-exposed surface of the RBD. ADI-55689, which potently neutralizes and competes with hACE2, appears to bind at the edge of the hACE2 binding site, close to the more structurally conserved core domain of the RBD, without overlapping with the CR3022 epitope (**Fig. 4B)**. ADI-56046, which exemplifies the group of antibodies that competes with both hACE2 and CR3022, binds slightly farther away from the flexible tip of the RBD and thus its epitope spans both the hACE2 binding site and the CR3022 epitope (**Fig 4C**). In summary, our structural analysis suggests that all of the nAbs recognize a single patch on the surface of the RBD with overlapping footprints. Given that these antibodies potently cross-neutralize SARS-CoV, SARS-CoV-2, and WIV1 suggests that this antigenic surface exhibits a high degree of conservation among the SARS-like coronaviruses.

**Figure 4.**
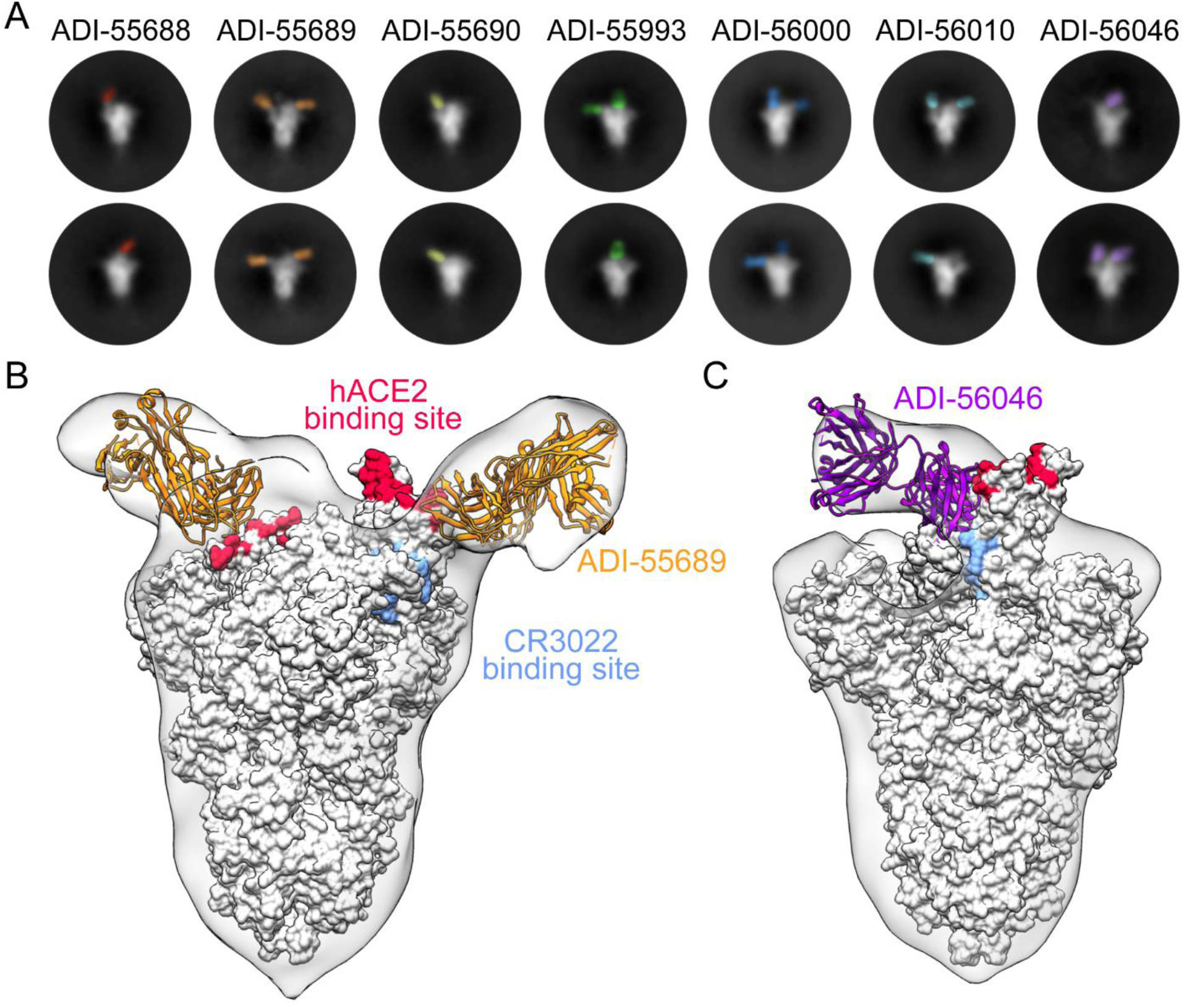
Structures of cross-neutralizing antibodies bound to SARS-CoV-2 S. (**A**) Negative-stain EM 2D class averages of SARS-CoV-2 S bound by Fabs of indicated antibodies. The Fabs have been pseudo-colored for ease of visualization. (**B-C**) 3D reconstructions of Fab:SARS-CoV-2 S complexes are shown in transparent surface representation (light gray) with the structure of the SARS-CoV-2 S trimer docked into the density (white surface). Fabs have been docked into the density and are shown in ribbon representation. S-bound Fabs of ADI-55689 (**B**) and ADI-56046 (**C**) are colored in orange and purple, respectively. The hACE2 and CR3022 binding sites on S are shaded in red and light blue, respectively.

In conclusion, we show that the cross-reactive human B cell response to SARS-CoV is composed of a broad diversity of clones that target multiple epitopes within the RBD, NTD, and S2 stem region. Unexpectedly, a large proportion of the antibodies in our panel displayed high levels of SHM, clonal expansion, and cross-reactivity with circulating HCoV S proteins, suggesting that SARS-CoV infection may have boosted a pre-existing MBC response induced by circulating HCoV exposure. This result is reminiscent of the phenomenon of “original antigenic sin” (OAS), whereby the primary viral exposure leaves an immunological imprint that shapes future immune responses to antigenically related viruses. OAS was first described in the context of repeated influenza exposures(*25, 26*), and more recently during secondary dengue infections(*27, 28*), but to our knowledge OAS has not been previously reported for CoVs. The OAS-phenotype antibodies described here show overall weak binding affinities for SARS-CoV and SARS-CoV-2 S and universally lack neutralizing activity against SARS-CoV-2, indicating that the corresponding serum antibody specificities are unlikely to mediate protection against these viruses or other SARS-like CoVs. Additionally, given previous reports suggesting the potential for antibody-mediated enhancement (ADE) of SARS-CoV infection in NHPs and feline infectious peritonitis virus in cats(*29, 30*), it will be important to investigate whether these non-neutralizing cross-reactive antibodies have any impact on SARS-CoV or SARS-CoV-2 pathogenesis. Future studies will be required to elucidate the role of pre-existing CoV immunity in modulating the antibody response to SARS-CoV-2 and the potential impact on disease outcome.

The potent cross-neutralizing antibodies described here bind to conserved epitopes overlapping the hACE2 binding site, thus illuminating this antigenic surface as a promising target for the rational design of pan-sarbecovirus vaccines. For example, the RBD epitope(s) defined by this class of antibodies could be presented on conformationally stable protein scaffolds to focus the antibody response on this site, as previously demonstrated for the motavizumab epitope on RSV F(*31*). Furthermore, the nAbs themselves, alone or in combination, represent promising candidates for prophylaxis or therapy of SARS, COVID-19, and potentially future diseases caused by new emerging SARS-like viruses.

## Acknowledgements

We thank E. Krauland and M. Vasquez for helpful comments on the manuscript. We also thank C. Kivler, C. O’Brien, E. Platt for antibody expression and purification, and M. Hagstroem, E. Worts and A. Gearhart for antibody sequencing. We acknowledge the generous provision of PBMCs from a SARS-1 survivor provided by Ingelise Gordon, Julie Ledgerwood, Wing-Pui Kong, Lingshu Wang, Kizzmekia Corbett, and other members of the NIAID Vaccine Research Center. This work was funded in part by a National Institutes of Health (NIH)/ National Institute of Allergy and Infectious Diseases (NIAID) grants awarded to J.S.M (R01-AI127521) and K.C. (R01-AI132633).

## References and notes

1. F. Wu et al., A new coronavirus associated with human respiratory disease in China. Nature 579, 265–269 (2020).

2. F. Li, Structure, Function, and Evolution of Coronavirus Spike Proteins. Annu Rev Virol 3, 237–261 (2016).

3. A. C. Walls et al., Cryo-electron microscopy structure of a coronavirus spike glycoprotein trimer. Nature 531, 114–117 (2016).

4. M. A. Tortorici, D. Veesler, Structural insights into coronavirus entry. Adv Virus Res 105, 93–116 (2019).

5. D. Wrapp et al., Cryo-EM structure of the 2019-nCoV spike in the prefusion conformation. Science 367, 1260–1263 (2020).

6. W. Song, M. Gui, X. Wang, Y. Xiang, Cryo-EM structure of the SARS coronavirus spike glycoprotein in complex with its host cell receptor ACE2. PLoS pathogens 14, e1007236 (2018).

7. J. Lan et al., Structure of the SARS-CoV-2 spike receptor-binding domain bound to the ACE2 receptor. Nature, (2020).

8. M. Hoffmann et al., SARS-CoV-2 Cell Entry Depends on ACE2 and TMPRSS2 and Is Blocked by a Clinically Proven Protease Inhibitor. Cell 181, 271–280 e278 (2020).

9. Q. Wang et al., Structural and Functional Basis of SARS-CoV-2 Entry by Using Human ACE2. Cell, (2020).

10. F. Li, Receptor recognition mechanisms of coronaviruses: a decade of structural studies. J Virol 89, 1954–1964 (2015).

11. S. Jiang, C. Hillyer, L. Du, Neutralizing Antibodies against SARS-CoV-2 and Other Human Coronaviruses. Trends Immunol, (2020).

12. X. Ou et al., Characterization of spike glycoprotein of SARS-CoV-2 on virus entry and its immune cross-reactivity with SARS-CoV. Nature communications 11, 1620 (2020).

13. F. Tang et al., Lack of peripheral memory B cell responses in recovered patients with severe acute respiratory syndrome: a six-year follow-up study. Journal of immunology (Baltimore, Md. : 1950) 186, 7264–7268 (2011).

14. M. S. Gilman et al., Rapid profiling of RSV antibody repertoires from the memory B cells of naturally infected adult donors. Science immunology 1, (2016).

15. T. F. Rogers et al., Zika virus activates de novo and cross-reactive memory B cell responses in dengue-experienced donors. Science immunology 2, (2017).

16. E. Goodwin et al., Infants Infected with Respiratory Syncytial Virus Generate Potent Neutralizing Antibodies that Lack Somatic Hypermutation. Immunity 48, 339–349 e335 (2018).

17. Z. A. Bornholdt et al., Isolation of potent neutralizing antibodies from a survivor of the 2014 Ebola virus outbreak. Science 351, 1078–1083 (2016).

18. A. Z. Wec et al., Longitudinal dynamics of the human B cell response to the yellow fever 17D vaccine. Proc Natl Acad Sci U S A 117, 6675–6685 (2020).

19. M. Yuan et al., A highly conserved cryptic epitope in the receptor-binding domains of SARS-CoV-2 and SARS-CoV. Science, (2020).

20. V. D. Menachery et al., SARS-like WIV1-CoV poised for human emergence. Proc Natl Acad Sci U S A 113, 3048–3053 (2016).

21. D. R. Burton, J. R. Mascola, Antibody responses to envelope glycoproteins in HIV-1 infection. Nat Immunol 16, 571–576 (2015).

22. A. C. Walls et al., Tectonic conformational changes of a coronavirus spike glycoprotein promote membrane fusion. Proc Natl Acad Sci U S A 114, 11157–11162 (2017).

23. A. C. Walls et al., Unexpected Receptor Functional Mimicry Elucidates Activation of Coronavirus Fusion. Cell 176, 1026–1039 e1015 (2019).

24. J. Pallesen et al., Immunogenicity and structures of a rationally designed prefusion MERS-CoV spike antigen. Proc Natl Acad Sci U S A 114, E7348–E7357 (2017).

25. G. Fazekas de St, R. G. Webster, Disquisitions of Original Antigenic Sin. I. Evidence in man. The Journal of experimental medicine 124, 331–345 (1966).

26. F. M. Davenport, A. V. Hennessy, T. Francis, Jr., Epidemiologic and immunologic significance of age distribution of antibody to antigenic variants of influenza virus. The Journal of experimental medicine 98, 641–656 (1953).

27. L. Priyamvada et al., B Cell Responses during Secondary Dengue Virus Infection Are Dominated by Highly Cross-Reactive, Memory-Derived Plasmablasts. J Virol 90, 5574–5585 (2016).

28. C. M. Midgley et al., An in-depth analysis of original antigenic sin in dengue virus infection. J Virol 85, 410–421 (2011).

29. H. Vennema et al., Early death after feline infectious peritonitis virus challenge due to recombinant vaccinia virus immunization. J Virol 64, 1407–1409 (1990).

30. Q. Wang et al., Immunodominant SARS Coronavirus Epitopes in Humans Elicited both Enhancing and Neutralizing Effects on Infection in Non-human Primates. ACS Infect Dis 2, 361–376 (2016).

31. B. E. Correia et al., Proof of principle for epitope-focused vaccine design. Nature 507, 201–206 (2014).

32. D. W. Zhang et al., RIP3, an energy metabolism regulator that switches TNF-induced cell death from apoptosis to necrosis. Science 325, 332–336 (2009).

33. L. M. Kleinfelter et al., Haploid Genetic Screen Reveals a Profound and Direct Dependence on Cholesterol for Hantavirus Membrane Fusion. mBio 6, e00801 (2015).

34. S. P. Whelan, L. A. Ball, J. N. Barr, G. T. Wertz, Efficient recovery of infectious vesicular stomatitis virus entirely from cDNA clones. Proc Natl Acad Sci U S A 92, 8388–8392 (1995).

35. M. Sarzotti-Kelsoe et al., Optimization and validation of the TZM-bl assay for standardized assessments of neutralizing antibodies against HIV-1. J Immunol Methods 409, 131–146 (2014).

36. T. Tiller et al., Efficient generation of monoclonal antibodies from single human B cells by single cell RT-PCR and expression vector cloning. J Immunol Methods 329, 112–124 (2008).

37. R. D. Gietz, R. A. Woods, Transformation of yeast by lithium acetate/single-stranded carrier DNA/polyethylene glycol method. Methods Enzymol 350, 87–96 (2002).

38. L. Shehata et al., Affinity Maturation Enhances Antibody Specificity but Compromises Conformational Stability. Cell reports 28, 3300–3308 e3304 (2019).

