## Supplementary Figures and Methods for "Broad sarbecovirus neutralizing antibodies define a key site of vulnerability on the SARS-CoV-2 spike protein" (from another version of this preprint: version 1, 2020-05-15)

**This PDF file includes:**

Materials and Methods
Supplementary Text
Figs. S1 to S12

### Materials and Methods

#### Human subjects.

Experimental guidelines of The U.S. Department of Health and Human Services were followed in the conduct of clinical research, and the Vaccine Research Center (VRC) protocol was approved by the National Institutes of Allergy and Infectious Diseases (NIAID), National institutes of Health (NIH) Institutional Review Board (IRB). Apheresis was performed on a survivor of the 2003 SARS outbreak after signed consent in accordance with NIH IRB reviewed protocols. Samples were processed to isolate peripheral blood mononuclear cells (PBMCs) and plasma in the laboratories of the VRC Vaccine Immunology Program. The isolated PBMCs and plasma were aliquoted and stored at -80°C until further use. The PBMC sample was sent to Adimab on March 19, 2020 for B cell isolation.

Human blood samples from healthy adult volunteers were collected after signed consent in accordance with IRB–reviewed protocols. Blood samples were processed in the Immune Monitoring Core Laboratory at the Dartmouth Geisel School of Medicine to isolate peripheral blood mononuclear cells (PBMCs) and plasma. The isolated PBMCs and plasma were aliquoted and stored at -80°C until further use. The healthy subjects were between the ages of 24-68 years at the time of blood-draw. None of the donors had a documented history of SARS-CoV or SARS-CoV-2 infection.

#### Cells.

Vero African grivet monkey kidney cells (obtained from ATCC) were passaged every 3 days using 0.05% Trypsin/EDTA solution (Gibco) and maintained in Dulbecco's Modified Eagle Medium (DMEM high glucose, Gibco) supplemented with 2% heat-inactivated fetal bovine serum (FBS, Atlanta Biologicals), 1% P/S (Gibco) and 1% Gluta-MAX (Gibco). HeLa and HEK293 cells were culture in DMEM (Invitrogen) supplemented with 10% fetal bovine serum (Invitrogen), 2 mM L-glutamine (Invitrogen), 100 units of Penicillin and 0.1 mg/ml of Streptomycin (Invitrogen). No antibiotics were included during lipofectamine transfections.

To generate HeLa-hACE2 reporter cells for the MLV pseudotype assays, hACE2 (NM_001371415) was cloned into pBOB(*32*) and co-transfected into HEK293T cells along with lentiviral packaging plasmids pMDL (Addgene), pREV (Addgene), and pVSVG (Addgene) using

Lipofectamine according to manufacturer's instruction in a 3:1:1:1 mass ratio. The media was exchanged after 16 hours. Supernatants containing hACE2 lentivirus were collected 32 hours later and stored at 4 degrees. 0.4 million HeLa cells were seeded per well of a 6-well plate. 24hrs later, 500 µl of hACE2-lentiviral supernatant was diluted with 1.5 ml of fresh media and added to cells with 10 µg/ml polybrene (Sigma) per well. The plate was centrifuged at 1500 x g for 30 minutes at room temperature. Fresh medium was added to the plate 12 h after transduction. Cell surface expression was confirmed by FACS using SARS-CoV-2 S-based probes.

HEK-293 cells (DSMZ, ACC 305) were suspension-adapted, cultured in CD OptiCHO AGT Medium (Gibco) supplemented with 6 mM GlutaMAX (Gibco) and maintained at 37°C in an orbital incubator shaker with humidified atmosphere containing 8% CO2. Vero C1008 (clone E6) cells (purchased from ATCC, CRL-1586) were cultured in Eagle's Minimum Essential Medium (EMEM) (ATCC, 30-2003) supplemented with 10% heat-inactivated fetal bovine serum (Avantor Seradigm FBS) and maintained at 37°C in a static incubator with humidified atmosphere containing 5% CO2.

**Generation of rVSV-SARS-CoV-2 and rVSV-SARS-CoV**

A recombinant vesicular stomatitis virus (rVSV) expressing an eGFP reporter and encoding the SARS-CoV-2 spike protein gene (GenBank: MN908947) in place of its native glycoprotein was generated by plasmid-based rescue as described previously(*33*, *34*). The identity of the generated virus was confirmed by Sanger sequencing of the spike protein encoding gene after RT-PCR amplification.

**Generation of MLV pseudovirus particles displaying SARS-CoV and SARS-CoV-2 S**

To generate pseudovirus, SARS-CoV (AAP13567) and SARS-CoV-2 (NC_045512) spike genes (codon optimized for mammalian expression) were synthesized (IDT) with 18 and 28 amino acid c-terminal deletions respectively and cloned into pCDNA3.3 (ThermoFisher). A luciferase reporter gene plasmid (addgene#18760) was modified to replace the IRES with a CMV promoter. These plasmids along with MLV gag/pol (addgene#14887) were purified using Endo-Free plasmid maxi kits (Qiagen). Under sterile conditions, these plasmids were co-transfected into HEK293T cells using Lipofectamine 2000 (ThermoFischer Scientific) according to the manufacturer's directions to produce single-round of infection competent pseudoviruses. 0.5 µg SARS-CoV-2 or

1 µg SARS-CoV-1 S, 2 µg gag/pol and 2 µg MLV luciferase was used per well of a 6-well plate. The media was exchanged 16 hours post-transfection. The supernatant containing SARS S-pseudotyped viral particles was collected 48h post transfection, aliquoted, and frozen at -80 °C for neutralization assays.

**Generation of authentic SARS-CoV-2**

Vero E6 cells were inoculated with SARS-CoV-2/MT020880.1 at an MOI = 0.01 and incubated at 37°C/5% $CO_2$/80% humidity. At 50 hours post-infection, cells were frozen at -80°C for 1 hour, allowed to thaw at room temperature, and supernatants were collected and clarified by centrifugation at ~2500xg for 10 minutes. Clarified supernatant was aliquoted and stored at -80°C. Sequencing data from this virus stock indicated a single mutation in the spike glycoprotein (H655Y) relative to Washington state isolate MT020880.1.

**Plaque Reduction Neutralization Assay, WIV-1**

Vero E6 cells (obtained from United States Army Medical Research Institute of Infectious Diseases) were seeded at $5x10^5$ cells/well in a 6-well plate 24h before the assay. One hundred plaque forming units (pfu) of infectious clone WIV-1 were mixed with antibody at 1:1 ratio and incubated at 37ºC for 30 min(*20*). Virus and Ab mix was added to each well and incubated for 45 min in a 37ºC + 5% C02 incubator. Plaque assay media (complete DMEM media + 0.8% low melting point agarose) was overlaid on top of the inoculum and incubated at 37ºC + 5% CO2 incubator for 48h. For plaque visualization, cells were stained with neutral red for at least 4 h before counting. Percent inhibition was calculated by: (# of plaques with no Ab – # of plaques)/# of plaques with no Ab x 100%. Fifty percent plaque reduction neutralization titer ($PRNT_{50}$) was calculated in GraphPad Prism 8.3.0 by fitting the data points using a sigmoidal dose-response (variable slope) curve.

**Recombinant VSV-SARS-CoV-2 neutralization assay**

A pre-titrated amount of rVSV-SARS2 S virus was incubated with serial dilutions of monoclonal antibodies or Fabs for 1 hr at room temperature. For screening the antibody-virus mixture was applied to monolayers of Huh7.5.1 cells in a 384-well plate and for titrations to monolayers of Vero cells in a 96-well plate. Following overnight incubation, eGFP-positive virus-infected cells were enumerated using a Cytation-5 imager (Biotek) and analyzed with the onboard Gen5 software.

**MLV pseudovirus neutralization assay**

SARS-CoV-1 and SARS-CoV-2 neutralization was assessed using a Murine Leukemia Pseudovirus assay. Briefly, SARS-CoV or SARS-CoV-2 S protein pseudotyped Murine Leukemia Virus (MLV) containing a firefly luciferase reporter gene was added to titrating amount of antibody and subsequently added to hACE2-expressing Hela cells where infection was monitored using a luminescence assay. The neutralization assay was performed as previously described with minor modifications(*35*). In sterile white 96-well half-area plates (Corning), 25µl of SARS-CoV or SARS-CoV-2 pseudotyped MLV vector was immediately mixed with 25 µl of serially diluted (5x using media) mAbs and incubated for one hour at 37°C to allow for antibody neutralization of the pseudotyped virus. 10,000 HeLa-hACE2 cells/well (in 50ul of media containing 20 µg/ml Dextran) were directly added to the antibody-virus mixture. Plates were incubated at 37°C for 42 to 48 h. Following the infection, HeLa-hACE2 cells were lysed using 1x luciferase lysis buffer (25mM Gly-Gly pH 7.8, 15mM $MgSO_4$, 4mM EGTA, 1% Triton X-100). Luciferase intensity was then read on a Luminometer with luciferase substrate according to the manufacturer's instructions (Promega, PR-E2620). Percentage of neutralization was calculated using the following equation.

$$100 * \left(1 - \frac{RULs\ of\ sample - Average\ RULs\ of\ Background}{Average\ of\ RULs\ of\ Virus\ only\ contrl - Average\ RULs\ of\ Backgroud}\right)$$

**Authentic SARS-CoV-2 neutralization assay**

A pre-titrated amount of authentic SARS-CoV-2 virus at final multiplicity of infection of 0.2. (Washington state strain MT020880.1) was incubated with serial dilutions of monoclonal antibodies for 1 h at room temperature. The antibody-virus mixture was applied to monolayers of Vero-E6 cells in a 96-well plate and incubated for 1 hour at 37°C in a humidified incubator. Infection media was then removed and cells were washed once with 1X PBS, followed by addition of fresh cell culture media. Culture media was removed 24 hours post infection and cells were washed once with 1X PBS. PBS was removed and plates were submerged in formalin fixing solution, then permeabilized with 0.2% Triton-X for 10 min at room temp and treated with blocking solution. Infected cells were detected using a primary detection antibody recognizing the SARS-CoV-2 nucleocapsid protein (Sino Biological) following staining with secondary detection antibody (goat α rabbit) conjugated to AlexaFluor 488. Infected cells were enumerated using Operetta high content imaging instrument and data analysis was performed using the Harmony software (Perkin Elmer).

**Production of recombinant SARS-CoV-1 and SARS-CoV-2 antigens**

Plasmids encoding the following constructs were transfected into FreeStyle 293F cells (Thermo Fisher) using polyethylenimine: residues 1-615 of human ACE2 with a C-terminal 8XHisTag and TwinStrepTag; residues 306-577 of the SARS-CoV-1 spike with a C-terminal monomeric Fc tag (SARS-CoV-1 RBD-SD1); residues 1-305 of the SARS-CoV-2 spike with a C-terminal monomeric Fc tag (SARS-CoV-2 NTD); residues 319-591 of the SARS-CoV-2 spike with a C-terminal monomeric Fc tag (SARS-CoV-2 RBD-SD1); residues 1-1276 of the HCoV-HKU1 spike with a "GSSGS" mutation from positions 752-756 and a C-terminal T4 fibritin motif, 8X HisTag and TwinStrepTag (HCoV-HKU1 S); residues 1-1276 of the HCoV-HKU1 spike with "PP" mutations at positions 1067 and 1068, "GSSGS" mutations from positions 752-756 and a C-terminal T4 fibritin motif, 8X HisTag and TwinStrepTag (HCoV-HKU1 S-2P); residues 1-1278 of the HCoV-OC43 spike with "PP" mutations at positions 1070 and 1071 and a C-terminal T4 fibritin motif, 8X HisTag and TwinStrepTag (HCoV-OC43 S-2P); residues 1-1190 of the SARS-CoV-1 spike with "PP" mutations at positions 968 and 969 a C-terminal T4 fibritin motif, 8X HisTag and TwinStrepTag (SARS-CoV-1 S-2P); residues 1-1208 of the SARS-CoV-2 spike with "PP" mutations at positions 986 and 987, "GSAS" mutations from positions 682-685 and a C-terminal T4 fibritin motif, 8X HisTag and TwinStrepTag (SARS-CoV-2 S-2P). Proteins were harvested from cell supernatants and constructs containing TwinStrepTags (ACE2, HCoV-HKU1 S, HCoV-HKU1 S-2P, HCoV-OC43 S-2P, SARS-CoV-1 S-2P and SARS-CoV-2 S-2P) were purified using StrepTactin resin (IBA), while constructs containing a monomeric Fc tag (SARS-CoV-1 RBD-SD1, SARS-CoV-2 NTD and SARS-CoV-2 RBD-SD1) were purified using Protein A resin (Pierce). HCoV-HKU1 S, HCoV-HKU1 S-2P, HCoV-OC43 S-2P, SARS-CoV-1 S-2P and SARS-CoV-2 S-2P were further purified by size exclusion chromatography using a Superose 6 Increase column (GE Healthcare) in 2 mM Tris pH 8.0, 200 mM NaCl and 0.02% $NaN_3$. ACE2, SARS-CoV-1 RBD-SD1, SARS-CoV-2 NTD and SARS-CoV-2 RBD-SD1were further purified by size exclusion chromatography using an S200 Increase column (GE Healthcare) in 2 mM Tris pH 8.0, 200 mM NaCl and 0.02% $NaN_3$.

### Commercially obtained human coronavirus S proteins

The S proteins from HCoV-229E, HCoV-NL63 and HCoV-OC43 were purchased from Sino Biological (Cat# 40605-V08B, 40604-V08B and 40607-V08B, respectively). The SARS-CoV-2 S1 subunit was purchased from Acro Biosystems (Cat# S1N-C52H3) and SARS-CoV-2 S2 subunit was purchased from Sino Biological (Cat# S2N-C52H5).

### Single B Cell sorting

B cells were purified from PBMCs using a MACS Human B Cells isolation kit (Miltenyi Biotec) and stained using anti-human CD19 (PE-Cy7; Biolegend), CD3 (PerCP-Cy5.5; Biolegend), CD8 (PerCP-Cy5.5; Biolegend), CD14 (PerCP-Cy5.5; Invitrogen), CD16 (PerCP-Cy5.5; Biolegend), propidium iodide, IgM (BV711; BD Biosciences), IgD (BV421; Biolegend), IgG (BV605; BD Biosciences), IgA (AF-488), CD27 (BV510; BD Biosciences), CD71 (APC-Cy7; Biolegend) and a mixture of PE- and APC-labelled SARS-CoV-2 S protein tetramers (25 nM each). Class-switched B cells, defined as $CD19^{+}CD3^{-}CD8^{-}CD14^{-}CD16^{-}IgM^{-}IgD^{-}$ cells, showing reactivity to both SARS-CoV-2 S tetramers were single-cell sorted using BD FACS Aria II (BD Biosciences) into 96-well polystyrene microplates (Corning) containing 20 µl /well of lysis buffer [5 µL of 5X first strand SSIV cDNA buffer (Invitrogen), 1.25 µL dithiothreitol (Invitrogen), 0.625 µL of NP-40 (Thermo Scientific), 0.25 µL RNaseOUT (Invitrogen), and 12.8 µL dH2O]. Plates were immediately stored at -80 ˚C. Flow cytometry data was analyzed using FlowJo software.

### Amplification and cloning of antibody variable genes

Human antibody variable gene transcripts (VH, Vκ, Vλ) were amplified by RT-PCR and nested PCRs using cocktails of IgM-, IgD-, IgA- and IgG-specific primers, as described previously(*36*). To allow cloning by homologous recombination into *S. cerevisiae,* the primers used in the second round of nested PCR contained 40 base pairs of 5' and 3' homology to the digested expression vectors. Amplified variable gene transcripts were transformed into yeast using the lithium acetate method for chemical transformation(*37*). Each transformation reaction contained 10 µL of unpurified VH and VL PCR product and 200 ng of the digested expression vectors. Following the transformation, individual yeast colonies were picked for sequencing and characterization.

**Expression and purification of IgGs**

IgGs used for binding experiments, competition assays, neutralization assays and structural studies were expressed in *S. cerevisiae* cultures grown in 24-well plates, as described previously(*14*). After 6 days of cell culture growth, the supernatant was harvested by centrifugation and IgGs were purified by protein A-affinity chromatography. The IgGs bound to the agarose were eluted with 200 mM acetic acid/50 mM NaCl (pH 3.5) into 1/8th volume 2 M Hepes (pH 8.0) and buffer-exchanged into PBS (pH 7.0).

The control antibody, CR3022, was cloned into *S. cerevisiae* using recombinational cloning. The variable region sequences of CR3022 were synthesized as gBlock fragments (IDT) with homologous overhangs which were cloned and produced as described above.

Fab fragments for structural studies were generated by digesting the IgGs with papain for 2 hours at 30 °C. The digestion was terminated by the addition of iodoacetamide, following which, the Fab and Fc mixtures were passed over Protein A agarose to remove Fc fragments and undigested IgG. The flow-through of the Protein A resin was then passed over CaptureSelect™ IgG-CH1 affinity resin (ThermoFisher Scientific). The Fabs captured on the resin surface were eluted with 200 mM acetic acid/50 mM NaCl (pH 3.5) into 1/8th volume 2 M Hepes (pH 8.0) followed by buffer exchange into PBS (pH 7.0).

**Bio-Layer Interferometry Kinetic Measurements (BLI).**

Apparent affinities ($K_D$) were measured by biolayer interferometry (BLI) using a FortéBio Octet HTX instrument (Molecular Devices). All steps were performed at 25 $^oC$ and at an orbital shaking speed of 1000 rpm. All reagents were formulated in PBSF buffer (PBS with 0.1% w/v BSA).

For experiments with IgGs immobilized on biosensors, the IgGs (100 nM) were captured (0.6 – 1.2 nm) onto anti-human IgG (AHC) biosensors (Molecular Devices) and then allowed to stand in PBSF for a minimum of 30 min. For experiments using Fc-fusion antigens (i.e. RBD-Fc or NTD-Fc), the IgG-loaded sensors were soaked (10 min) in an irrelevant IgG1 solution (0.5 mg/mL) to block remaining Fc binding sites. For experiments using strep-tagged antigens, the IgG-loaded sensors were soaked (10 min) in a biocytin solution (100 uM) to saturate remaining streptavidin binding sites. After a short (60 s) baseline step in PBSF, the IgG-loaded biosensors were exposed (180 s) to the antigen (100 nM) and then dipped (180 s) into PBSF to measure any dissociation of the antigen from the biosensor surface. Data for which binding responses were >0.1 nm were aligned, inter-step corrected (to the association step) and fit to a 1:1 binding model using the FortéBio Data Analysis Software, version 11.1.

Experiments with IgGs or Fabs in solution were performed using biotinylated or Fc-fusion proteins. Recombinant biotinylated antigens (100 nM) were immobilized (1.0 – 2.0 nm) on streptavidin biosensors (Molecular Devices). Fc-fusion antigens (100 nM) were captured (1.0 – 2.0 nm) to AHC biosensors (Molecular Devices), and then soaked (10 min) in an irrelevant IgG1 solution (0.5 mg/mL) to block remaining Fc binding sites (only for IgG in solution experiments). All sensors were then allowed to stand in PBSF for a minimum of 30 min. After a short (60 s) baseline step in PBSF, the antigen-loaded biosensors were exposed (180 s) to the IgG or Fab (100 nM) and then dipped (180 s) into PBSF to measure any dissociation of the IgGs from the biosensor surface. Data for which binding responses were >0.1 nm were aligned, interstep corrected (to the association step) and fit to a 1:1 binding model using the FortéBio Data Analysis Software, version 11.1.

**Competition analysis using biolayer interferometry (BLI)**

Competition of antibodies with CR3022 and hACE2 for binding to recombinant SARS-CoV-2 RBD was evaluated using a ForteBio Octet HTX instrument (Molecular Devices). All binding steps were performed at 25$^{o}$C and at an orbital shaking speed of 1000 rpm. All reagents were formulated in PBSF buffer (PBS with 0.1% w/v BSA). The IgGs (100 nM) were captured to anti-human IgG capture (AHC) biosensors (Molecular Devices) to a sensor response of 1.0nm-1.4 nm, the remaining unoccupied binding sites on the biosensor were blocked with an inert IgG (0.5mg/mL), and then allowed to equilibrate in PBSF for a minimum of 30 min. To assess any cross interactions between proteins on the sensor surface and the secondary molecules, the loaded and blocked sensors were exposed (90 s) to hACE2 receptor (300 nM) or CR3032 IgG (300 nM) prior to the binning analysis. After a short baseline step (60 s) in PBSF, the IgG-loaded biosensor tips were exposed (180 s) to the SARS-CoV-2 RBD-SD1 antigen (100 nM) and then exposed (180 s) to either the hACE2 soluble receptor (300 nM) or CR3022 IgG (300 nM). The data was y-axis normalized, and interstep corrected using the FortéBio Data Analysis Software version 11.0. Additional binding by the secondary molecule indicates an unoccupied epitope (non-competitor), while no binding indicates epitope blocking (competitor).

**Surface plasmon resonance Fab affinity determination**

Tagged SARS-CoV-2 S-2P was immobilized to a single flow cell of a NiNTA sensorchip (GE Healthcare) at a level of ~700 response units (RUs) in a Biacore X100 (GE Healthcare). Fabs, diluted in SPR running buffer composed of 10 mM HEPES pH 8.0, 150 mM NaCl and 0.05% Tween 20, were injected over both flow cells at two-fold serial dilutions with concentrations ranging from 100-6.25 nM. The sensorchip was doubly regenerated using 350 mM EDTA pH 8.3 and 100 mM NaOH. Data were double-reference subtracted and fit to a 1:1 binding model using Biacore X100 Evaluation Software (GE Healthcare).

**Serum ELISAs**

For analysis of human serum reactivity to HCoV S proteins, 96-well polystyrene ELISA plates (Corning) were coated with 25 μl per well of S protein at a concentration of 5 μg/ml diluted in 1X PBS and incubated overnight at 4 °C. Wells were washed 3 times with 1X PBS and then blocked with 5% non-fat dried milk (NFDM) in 1X PBS for 1 h at 37 °C. Wells were washed and then blocked with 5% non-fat dried milk (NFDM) in PBS for 1 h at 37 °C. Wells were washed three times with PBS and serial dilutions of human plasm in 5% NFDM-PBS were added and incubated for 1 h at 37 °C. Plates were then washed three times with PBS and secondary cross-adsorbed the anti-human IgG-HRP (Thermo Fisher Scientific) detection antibody was added at 1:8000 dilution in 5% NFDM-PBS for 1 h at 37 °C. After washing three times with PBS, 25 ul per well of the 1-Step™ Ultra TMB-ELISA Substrate Solution (Thermo Fisher Scientific) was added to detect binding as per manufacturer recommendations, followed by addition of an equal volume of the stop solution and absorbance was measured at 450 nm wavelength using a Spectramax microplate Reader (Molecular Devices).

**Polyreactivity Assay**

Antibody polyreactivity was measured as described previously(*38*). Briefly, soluble membrane protein (SMP) and soluble cytosolic protein (SCP) fractions obtained from Chinese hamster ovary (CHO) cells were biotinylated using NHS-LC-Biot (Thermo Fisher Scientific). IgGs presented on the surface of yeast were incubated with 1:10 diluted biotinylated CHO cell preparations on ice for 20 minutes. Cells were then washed twice with ice-cold PBS containing 0.1% BSA (PBSF) and incubated in 50 μL of a secondary labelling mix containing ExtrAvidin-R-PE (Sigma-Aldrich), anti-human LC-FITC (Southern Biotech) and propidium iodide) for 15 minutes. The cells were washed twice with PBSF and resuspended in PBSF to be run on a FACSCanto II (BD Biosciences). The mean fluorescence intensity of binding was normalized using control antibodies that display low, medium or high polyreactivity to assess the non-specific binding.

**Cell surface-expression of recombinant SARS-CoV and SARS-CoV-2 S proteins and evaluation of antibody binding**

Suspension-adapted HEK-293 cells were grown to a density of 4 x $10^6$ cells/mL in CD OptiCHO AGT Medium (Gibco) supplemented with 6 mM GlutaMAX (Gibco). For transient transfection, HEK-293 cells were pelleted and resuspended in FreeStyle F17 Expression Medium (Gibco) supplemented with 6 mM GlutaMAX. DNA plasmids encoding the full length wild type SARS-CoV-2 or SARS-CoV-2P spike proteins or the empty vector plasmid (260 ug) were incubated with PEIpro (525 uL) (PolyPlus) in FreeStyle F17 Expression Medium with 6 mM GlutaMAX for 10-15 minutes at room temperature before addition to the HEK-293 cell suspension (final cell density of 4 x $10^6$ cells/mL in 200mL volume). Transfected cultures were maintained at 37°C in an orbital incubator shaker with humidified atmosphere containing 8% $CO_2$ and harvested 24 or 48 hours after transfection. To verify cell surface expression, transfected cells were labeled with antibody CR3022 as described below.

Transiently transfected cells and mock transfected cells were washed with cold PBSF and 200,000 cells per well were then aliquoted into 96 well plates. Cells were resuspended in 100 uL of each test monoclonal antibody at 100 nM and incubated on ice for 30 minutes. Afterwards, cells were washed two times with cold PBSF and resuspended in 50 uL of secondary stain containing goat anti-human IgG PE at a 1:100 dilution (SouthernBiotech) and propidium iodide and incubated on ice and in the dark for 20 minutes. After completion of the secondary stain, cells were washed twice with cold PBSF and resuspended with 50 uL of fresh PBSF. Cell binding was analyzed by flow cytometry using a BD FACSCanto II (BD Biosciences) equipped with a 96 well plate high throughput sampler. Binding was determined by calculating the fold over background binding ratio: MFI values of target expressing cells divided by the MFI of mock transfected cells.

On-cell, twelve- or eight- point titrations of selected monoclonal antibodies were performed as described above on WT SARS-CoV-2 S expressing cells over a three- or four- fold dilution series respectively, starting at 300 nM concentration. Percent binding was determined by subtracting the MFI value of an isotype control from the test antibodies and normalizing to the maximum MFI value for each antibody: ($MFI_{mAb}$ – $MFI_{control}$) / Max $MFI_{mAb}$ x 100. $EC_{50}$ values were calculated by non-linear regression (sigmoidal) using GraphPad Prism 8.2.1 software.

**Antibody binding to cell surface expressed SARS-CoV-2 S over time**

HEK293 cells expressing cell-surface WT SARS-CoV-2 S were generated as described above. 200,000 cells in 60 uL of FreeStyle F17 medium were added to a 96-well plate. 300 nM antibody in PBSF and the cells were equilibrated separately for 15 minutes to 37 °C. At each timepoint (240, 180, 120, 60, 30, 20, 10, 5 minutes), 30 uL of each antibody were mixed with the cell suspension to a final concentration of 100 nM antibody. After incubation for the allotted time, the samples were immediately placed on ice and washed twice with ice-cold PBSF before staining cells with 50 uL of secondary stain containing goat anti-human IgG PE at a 1:100 dilution and propidium iodide. After 20 minutes on ice in the dark, cells were washed twice with PBSF and resuspended with 50 uL of fresh PBSF and analyzed by flow cytometry using a BD FACSCanto II (BD Biosciences) equipped with a 96 well plate high throughput sampler. Binding was determined at each timepoint by the MFI and each sample was normalized to the MFI at the 5 minute timepoint (MFI / $MFI_{5\ min}$ x 100).

**Antibody inhibition of SARS-CoV-2 S binding to cell-surface ACE2**

Monoclonal antibody and SARS-CoV-2 S-2P were mixed to a final concentration of 500 nM antibody and 100 nM SARS-CoV-2 S-2P in PBSF and incubated on ice for 20 minutes. 50 uL of the mixture was used to resuspend 200,000 PBSF-washed Vero E6 cells cells (VERO C1008 [Vero 76, clone E6, Vero E6]; ATCC No. CRL-1586) and incubated on ice for 20 minutes. The cells were then washed twice with PBSF and stained with Strep-Tactin PE (IBA Lifesciences; Cat.no: 6-5000-001) at a 1:200 dilution and propidium iodide. Cells were washed twice with cold PBSF before being resuspended with 50 uL of fresh PBSF and analyzed by flow cytometry using a BD FACSCanto II (BD Biosciences) equipped with a 96 well plate high throughput sampler. Percent SARS-CoV-2 S-2P binding was calculated by: (% positive cells with test antibody / % positive cells with isotype control) x 100.

**Negative stain electron microscopy**

SARS-CoV-2 S-2P was diluted to a concentration of 0.04 mg/mL using 2 mM Tris pH 8.0, 200 mM NaCl and 0.02% $NaN_3$ before being mixed with a 5-fold molar excess of Fab. Fab was allowed to bind on ice for 5 minutes before the mixture was deposited on plasma cleaned CF-400 grids (EMS) and stained using methylamine tungstate (Nanoprobes). Grids were imaged at a nominal magnification of 92,000X (corresponding to a calibrated pixel size of 1.63 Å/pix) in a Talos F200C TEM microscope equipped with a Ceta 16M detector (ThermoFischer Scientific). CTF estimation and particle picking were performed in cisTEM (Grant et al, eLife 2018) and extracted particles were exported to cryoSPARC v2 (Punjani et al, Nature Methods 2017) for 2D classification and *ab initio* 3D reconstruction and heterogenous refinement.

**Supplementary Figures**

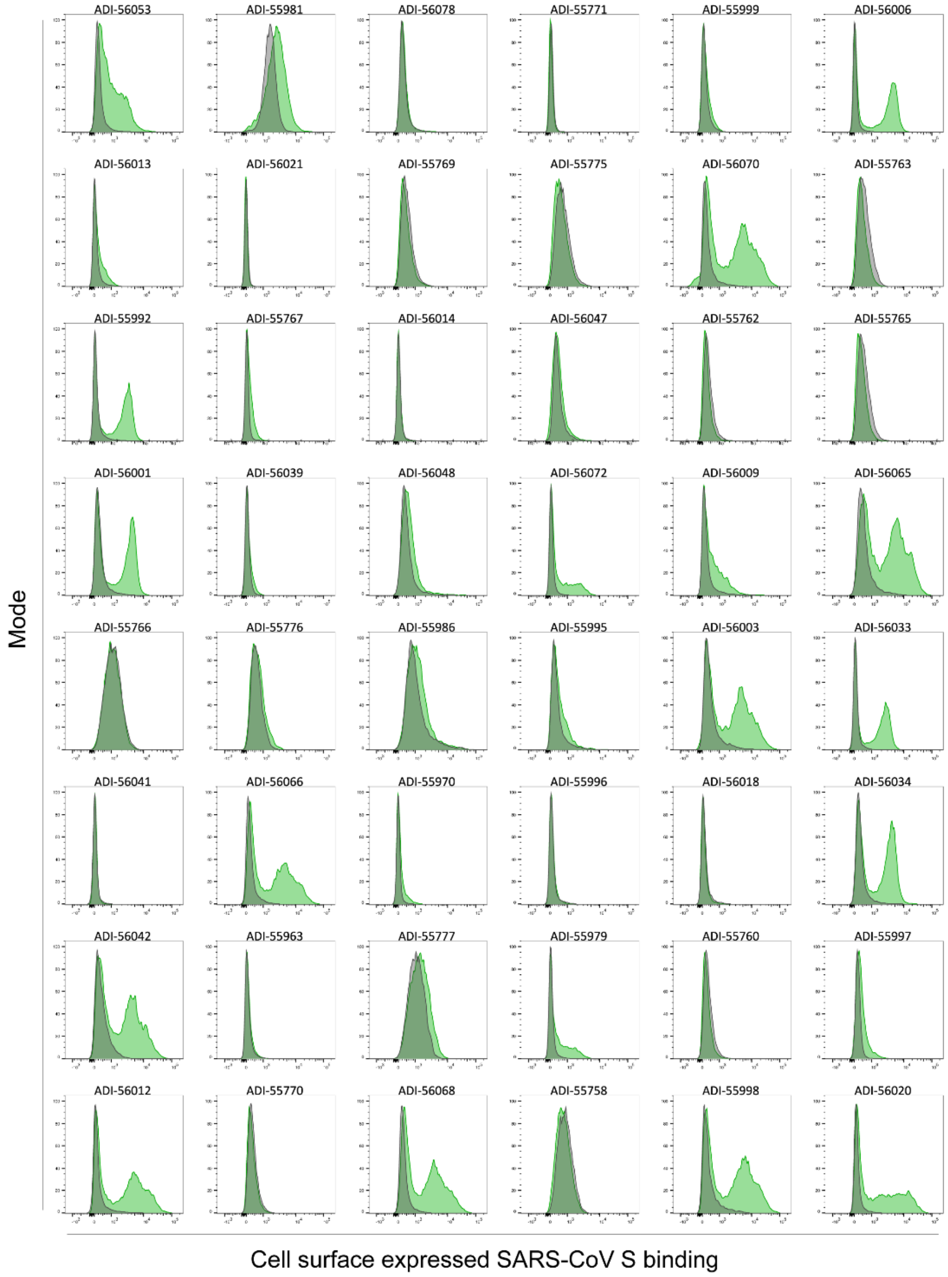


**Fig. S1.** Binding of SARS-CoV-2 S-specific antibodies to SARS-CoV S expressed on the surface of HEK293 cells. Antibodies were tested at a 100 nM concentration. The antibodies shown did not show detectable binding to recombinant SARS-CoV S, as determined by BLI. Green trace indicates cell surface binding of test IgGs to HEK293 cells transfected with SARS-CoV S; grey traces show test IgG binding to empty-vector transfected HEK293 cells.

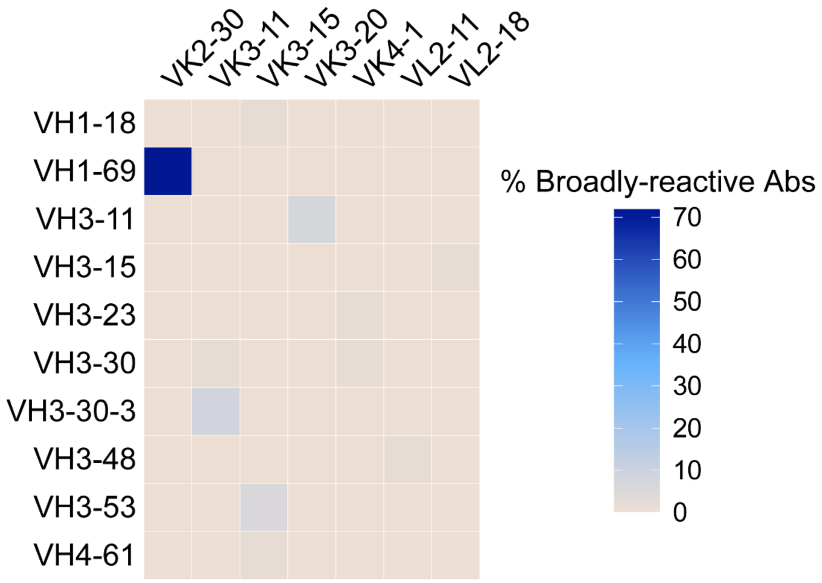


**Fig. S2.** $V_H/V_L$ germline gene usage of SARS-CoV-2 S-reactive antibodies that display cross-reactivity with one or more circulating HCoV S protein.

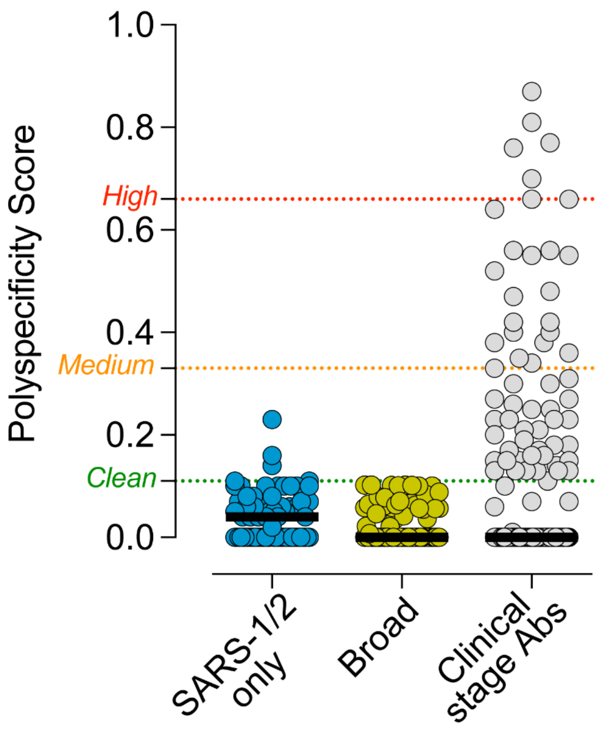


**Fig. S3.** Polyreactivity of broadly cross-reactive and SARS-CoV/SARS-CoV-2-specific antibodies, as determined using a previously described assay(*31*). Thresholds for high, medium, low, and no polyreactivity are indicated by dashed lines. Polyspecificity scores for 138 clinical stage antibodies are shown for comparison. Type or paste caption here. Create a page break and paste in the Figure above the caption.

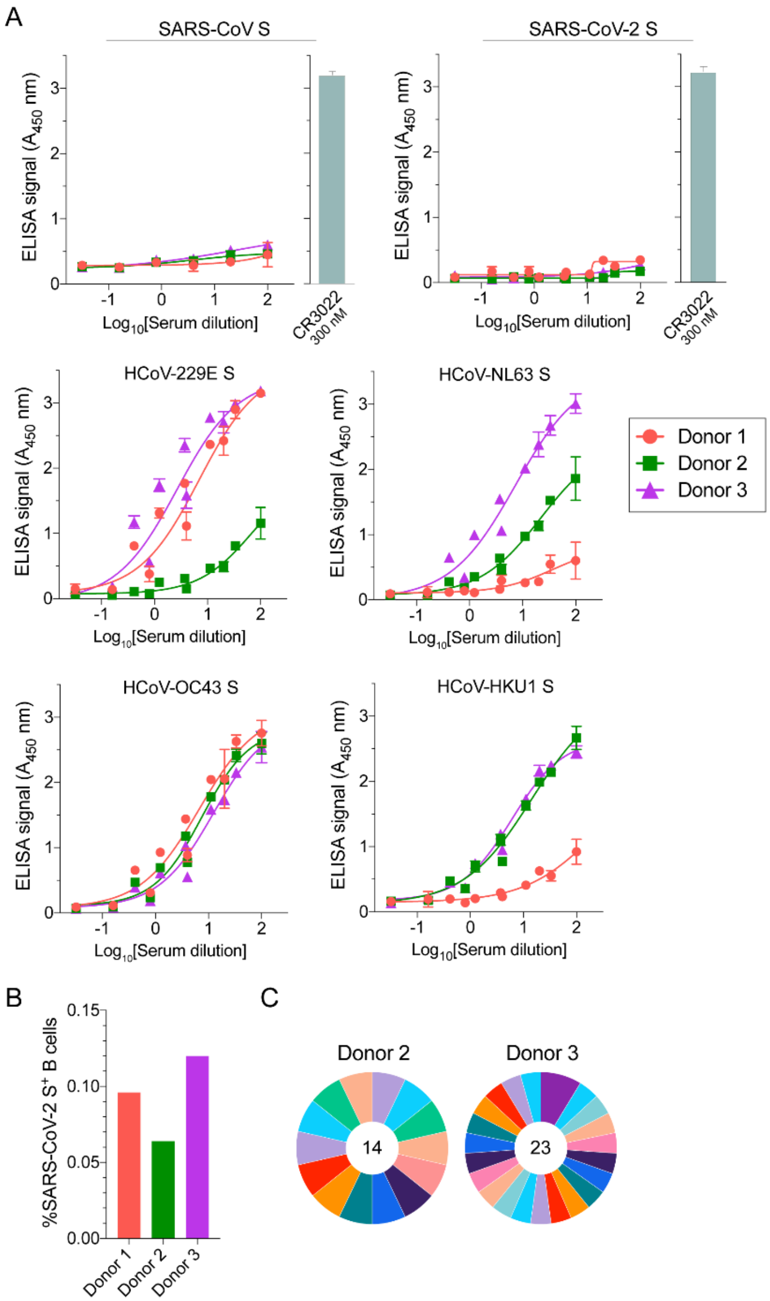


S6

**Fig. S4.** (A) ELISA binding reactivity of healthy donor sera to SARS-CoV, SARS-CoV-2, and circulating HCoV S proteins. CR3022 is included as a positive control. (B) Percentage of SARS-CoV-2 S-specific B cells observed in each naïve donor, as determined by flow cytometry. The percentages shown indicate the frequency of SARS-CoV-2 S-reactive B cells among $CD19^+$ B cells. (C) Clonal lineage analysis. Each lineage is represented as a segment proportional to the lineage size. The total number of isolated antibodies is shown in the center of each pie. Clonal lineages were defined based on the following criteria: identical VH and VL germline gene, identical CDR H3 length, and CDR H3 amino acid identity ≥80%. Clonal lineages are not shown for Donor 1 because only three SARS-CoV-2 S-specific antibodies were isolated.

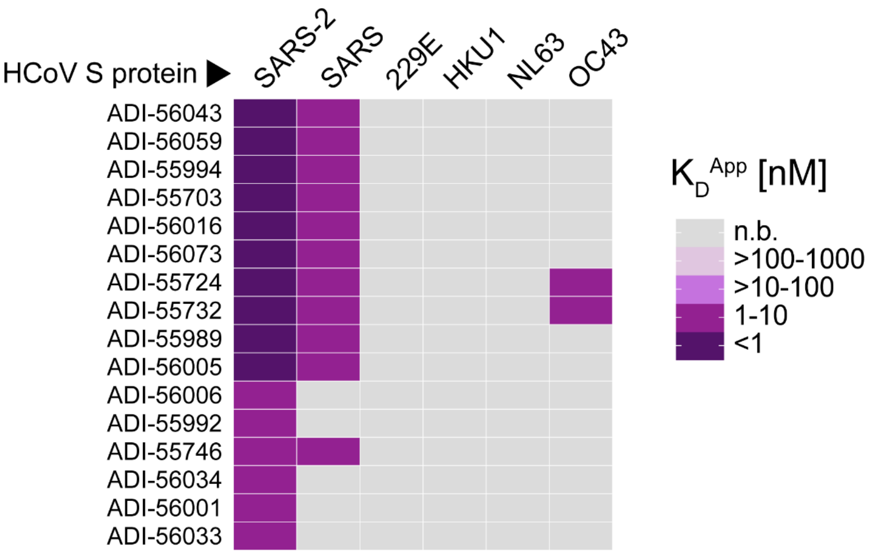


**Fig. S5.** Heat map showing $K_D^{Apps}$ of SARS-CoV-2 S2-specific antibodies for SARS-CoV, HCoV-HKU1, HCoV-229E, HCoV-OC43, and HCoV-NL63 S proteins, as determined by BLI. N.B., non-binder.

| Clone name | CR3022 competition trace | CR3022 competitor? | hACE2 competition trace | hACE2 competitor? |
|---|---|---|---|---|
| ADI-55689 | 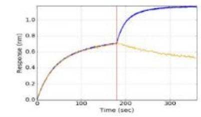 | No | 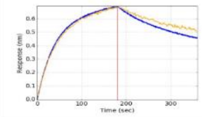 | Yes |
| ADI-55699 | 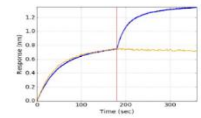 | No | 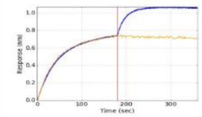 | No |
| ADI-55754 | 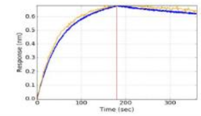 | Yes | 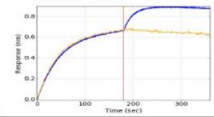 | No |
| ADI-56051 | 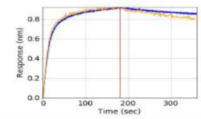 | Yes | 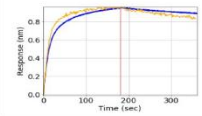 | Yes |

**Fig. S6.** Representative sandwich binning sensorgrams to competitor proteins, CR3022 and hACE2.

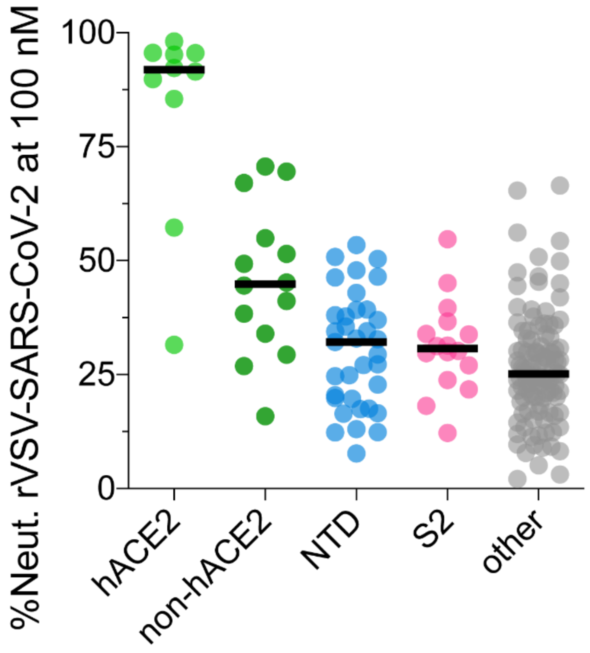


**Fig. S7.** Single-point neutralization screening of mAbs with a VSV-SARS-CoV-2 pseudovirus assay. The neutralizing activity of the entire collection of mAbs, each at 100 nM, was determined using Vero target cells. Antibodies are grouped according to epitope specificity. hACE2, RBD-directed antibodies that compete with hACE2; non-hACE2, RBD-directed antibodies that do not compete with hACE2. Black bars indicate medians.

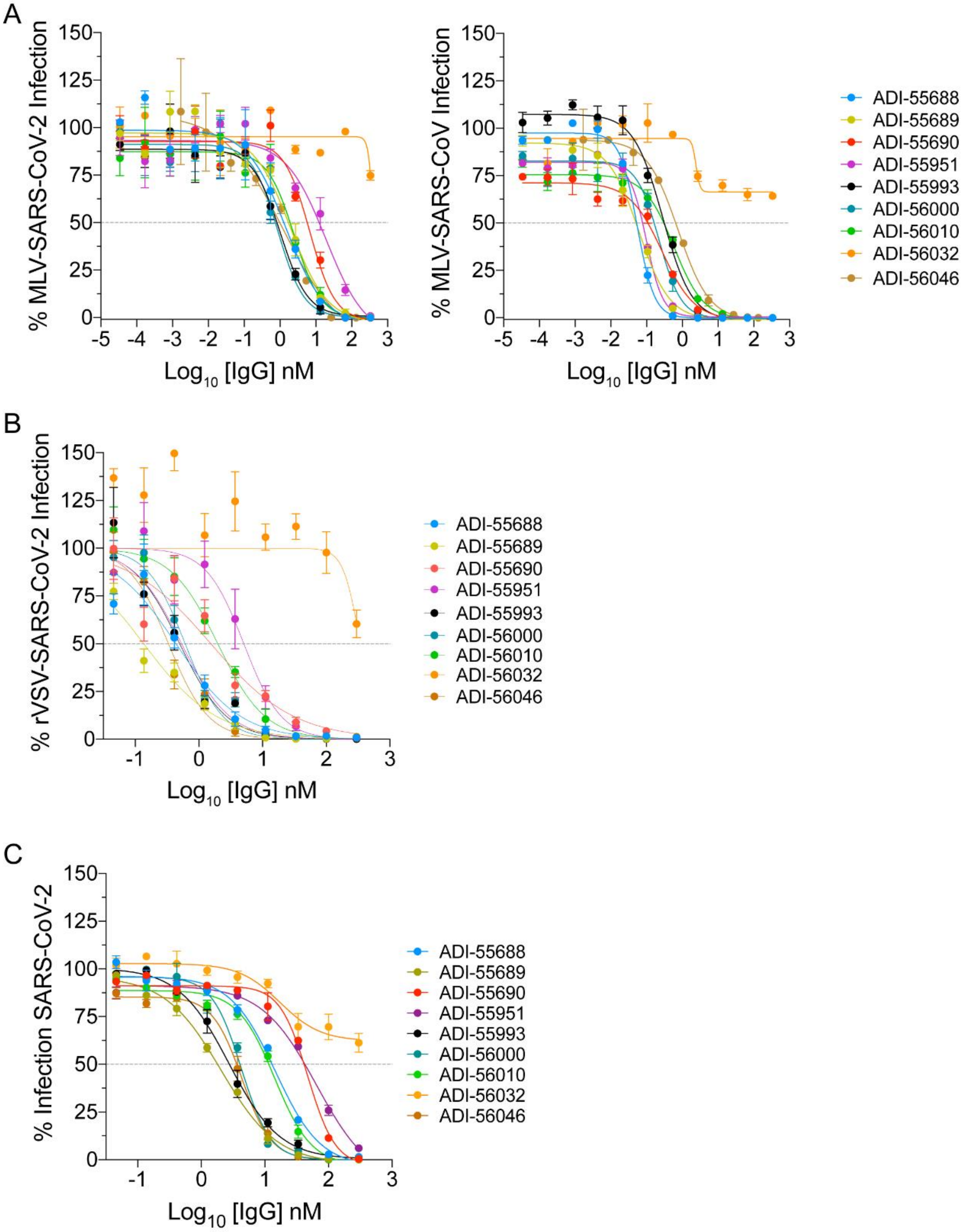


**Fig. S8.** Representative SARS-CoV-2 neutralization curves. Neutralization titration experiments were performed using an MLV-SARS-CoV-2 pseudovirus assay using HeLa-ACE2 target cells (A), a VSV-SARS-CoV-2 pseudovirus assay using Vero target cells and (C) an authentic virus assay using Vero E6 target cells. The curves were fitted for nonlinear regression (log [inhibitor] vs. normalized response, variable slope).

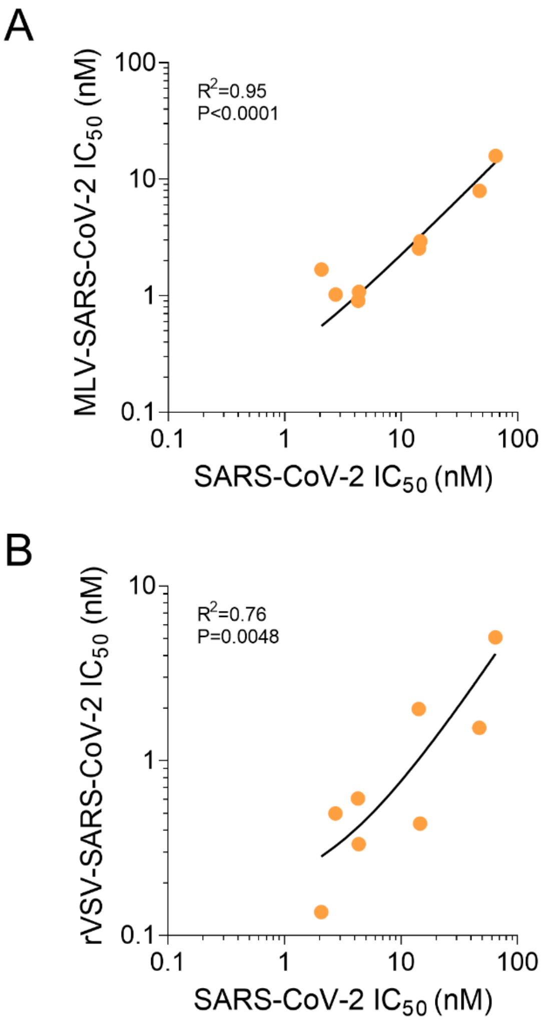


**Fig. S9.** Correlation between authentic SARS-CoV-2 neutralization $IC_{50}$s using Vero E6 target cells and (A) MLV-SARS-CoV-2 pseudovirus neutralization $IC_{50}$s using HeLa-ACE2 target cells or (B) VSV-SARS-CoV-2 neutralization $IC_{50}$s using Vero E6 target cells. $R^2$ values were calculated by linear regression.

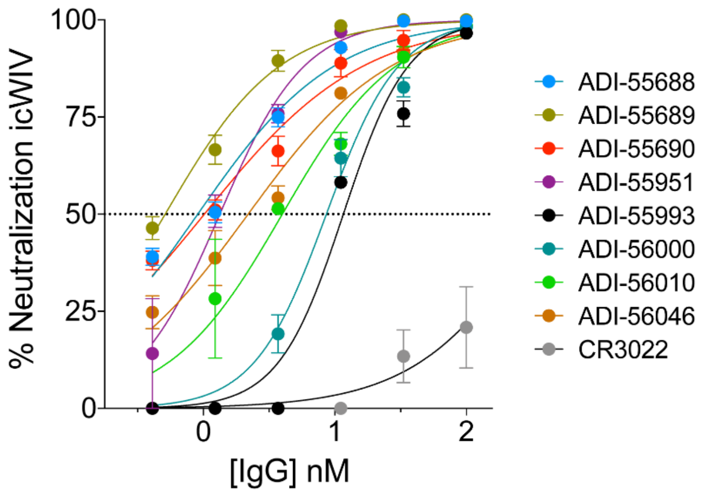


**Figure S10.** WIV1 neutralization curves. Neutralization titration experiments were performed using a WIV1 plaque reduction assay using Vero E6 target cells. The curves were fitted by nonlinear regression [sigmoidal dose-response (variable slope)]. CR3022 is included for comparison. ADI-26140 is negative control antibody specific for hen egg lysozyme.

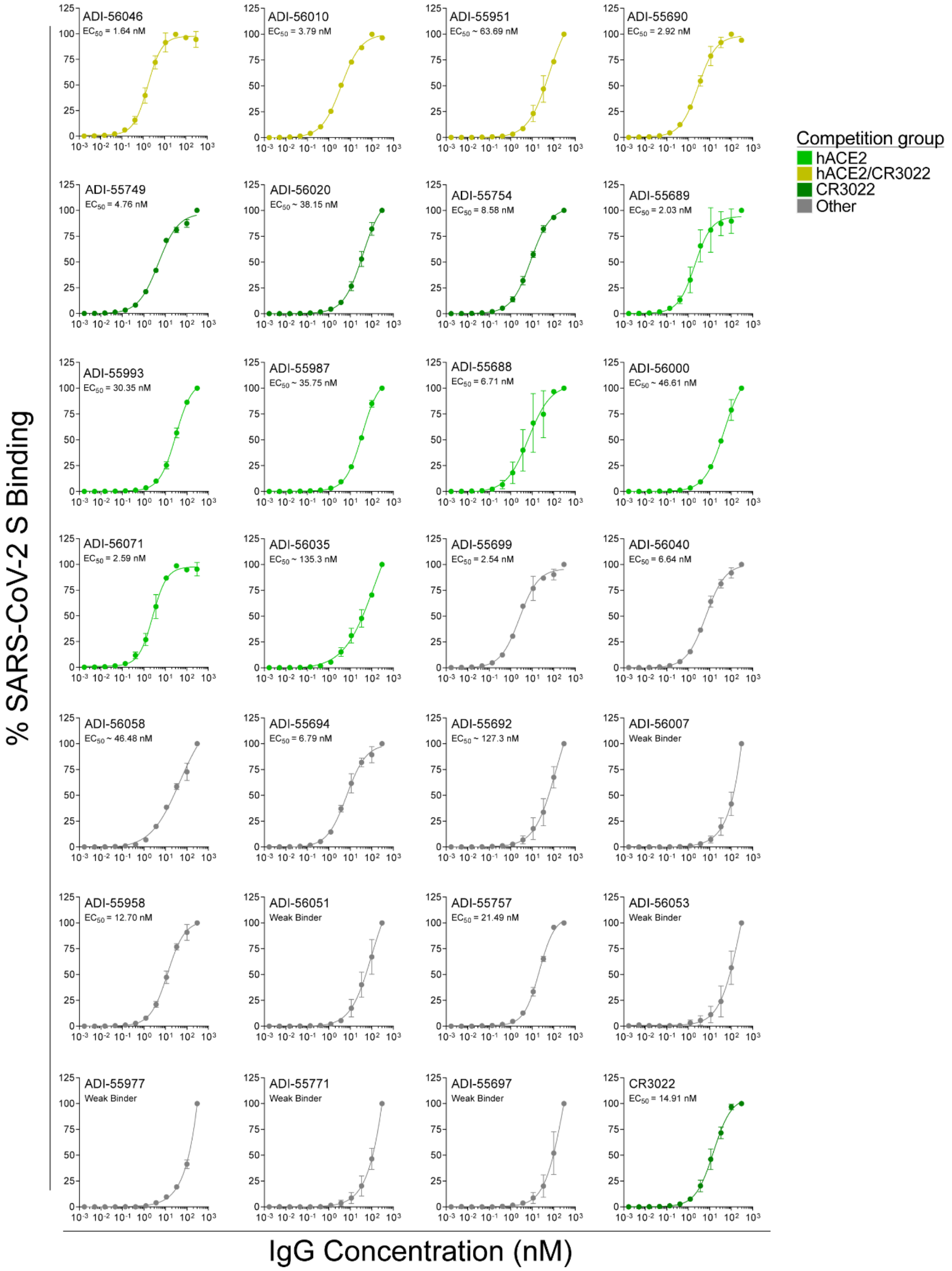


**Figure S11.** Binding of RBD-directed antibodies to WT SARS-CoV-2 S expressed on the surface of HEK293 cells. Antibody binding was detected using an anti-human IgG-PE secondary antibody. Curves are colored based on epitope specificity: hACE2, antibodies that only compete with hACE2; CR3022, antibodies that only compete with CR3022; hACE2/CR3022, antibodies that compete with hACE2 and CR3022; Other, antibodies that do not compete with hACE2 or CR3022. Binding $EC_{50}$s were calculated using a non-linear regression analysis.

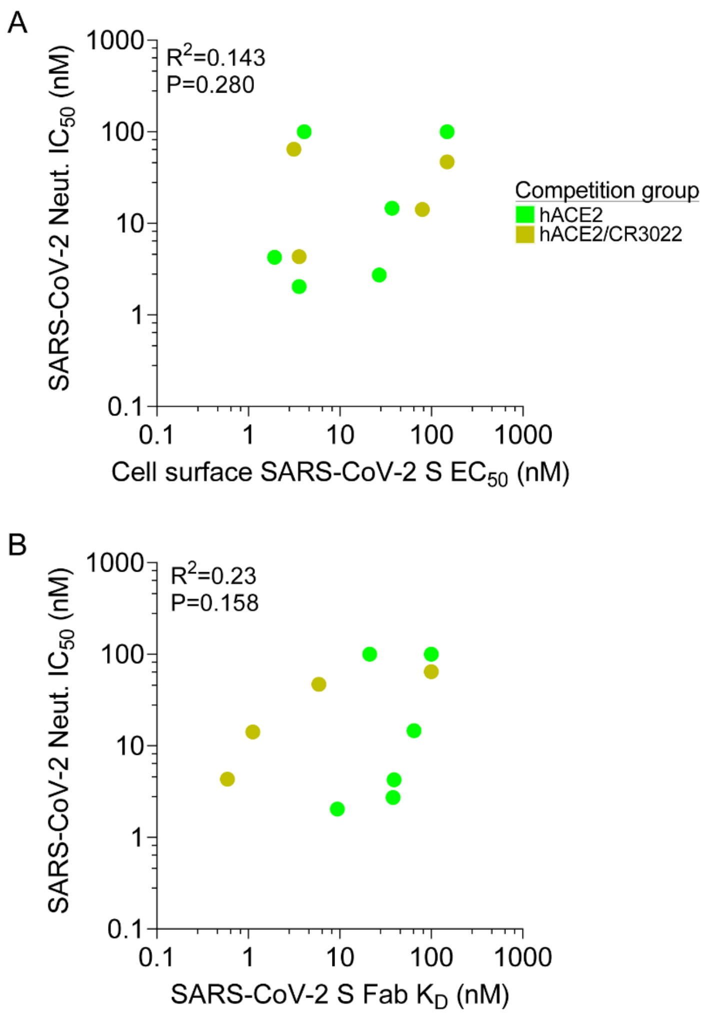


**Figure S12.** (A) Authentic SARS-CoV-2 neutralization $IC_{50}$s are plotted against binding half maximal effective concentrations ($EC_{50}$s) for SARS-CoV-2 S expressed on the surface of cells. (B) Authentic SARS-CoV-2 neutralization $IC_{50}$s are plotted against Fab binding affinities for recombinant SARS-CoV-2 S. Only hACE2 blocking antibodies are shown. $R^2$ and p values are indicated.
